# Image analysis for taxonomic identification of Javanese butterflies

**DOI:** 10.1101/408146

**Authors:** Saskia de Vetter, Rutger Vos

## Abstract

Taxonomic experts classify millions of specimens, but this is very time-consuming and therefore expensive. Image analysis is a way to automate identification and was previously done at Naturalis Biodiversity Center for slipper orchids (Cypripedioideae) by the program ‘OrchID’. This program operated by extracting a pre-defined number of features from images, and these features were used to train artificial neural networks (ANN) to classify out-of-sample images. This program was extended to work for a collection of Javanese butterflies, donated to Naturalis by the Van Groenendael-Krijger Foundation. Originally, for the orchids, an image was divided into a pre-defined number of horizontal and vertical bins and the mean blue-green-red values of each bin were calculated (BGR method) to obtain image features. In the extended implementation, characteristic image features were extracted using the SURF algorithm implemented in OpenCV and clustered with the BagOfWords method (SURF-BOW method). In addition, a combination of BGR- and SURF-BOW was implemented to extract both types of features in a single dataset (BGR-SURF method). A selection of the butterfly and orchid images was made to create datasets with at least 5 and at most 50 specimens per species. The SURF-BOW and BGR-SURF methods were applied to both selected datasets, and the original BGR method was applied to the selected butterfly dataset. PCA plots were made to inspect visually how well the applied methods discriminated among the species. For the butterflies, both genus and species appeared to cluster together in the plots of the SURF-BOW method. However, no obvious clustering was noticeable for the orchid plots. The performance of the ANNs was validated by a stratified *k*-fold cross validation. For the butterflies, the BGR-SURF method scored best with an accuracy of 77%, against 71% for the SURF-BOW method and 66% for the BGR method, all for chained genus and species prediction with *k* = 10. The new methods could not improve the accuracy of the orchid classification with *k* = 10, which was 75% on genus, 52% on genus and section and 48% on genus, section and species in the original framework and now less than 25% for all. The validation results also showed that at least about 15 specimens per species were necessary for a good prediction with the SURF-BOW method. The BGR-SURF method was found to be the best of these methods for butterflies, but the original BGR method was best for the slipper orchids. In the future these methods may be tested with other datasets, for example with mosquitoes. In addition, other classifiers may be tested for better performance, like support vector machines.

## Introduction

Ever since Carolus Linnaeus proposed his hierarchical classification system in the 1700s, scientists have been classifying species and higher taxa to structure life on Earth [1]. Experts in taxonomy have classified millions of specimens by hand, but this is very time-consuming and therefore expensive, and as not every collected specimen is named yet and more and more are collected, there is a continuing need for more scalable taxonomic identification [2].

A solution to this problem is to use image analysis to recognize species and to classify them automatically to the lowest possible level of taxonomy. This has been done before, for example for slipper orchids (Cypripedioideae), by making use of artificial neural networks trained on image features [3]. Pictures of these orchids that were already classified were used as a reference to train neural networks. Then, out-of-sample pictures of orchids that belong to this group could be identified based on their image features. This resulted in the development of ‘OrchID’, a framework to identify slipper orchids by image analysis [3].

Naturalis Biodiversity Center [4] recently received a large collection of Javanese butterflies, collected in the 1930s and since held by the Van Groenendael-Krijger Foundation, which has not been fully identified yet. At the time, each butterfly was put in a little envelope (a “papillotte”), which have since been stored in drawers. The goal of this project is to add more features to the existing OrchID framework in order to classify these Javanese butterflies and to make the recognition of the slipper orchids, if possible, more specific.

### Image data and metadata management

Prior to this project, pictures of some of the butterflies in the collection had been taken in standard orientation and with a ruler next to it. These pictures were uploaded to the photo hosting website Flickr (www.flickr.com) on an account associated with the OrchID framework. On this account, the pictures of the slipper orchids were stored as well. Since the butterfly collection is very large (the total number of specimens is not known but runs in the thousands), pictures of other specimens were continuously being taken during this project and progressively became and continue to become available to use in the dataset. Every time a new part of the collection becomes available (hundreds at a time), this new dataset is processed by the program to test its functionalities and to see if fine-tuning of the framework is necessary.

The Flickr website provides the ability to add ‘tags’ to a picture, which means that metadata can be added to a picture to provide information on what the image is about. In the case of the butterflies, the tags are the different hierarchical ranks as far as the taxonomy is known for that picture. The information for the tags comes from butterfly taxonomist Jan Moonen, who identified part of this butterfly collection so it could be used as training data. When the identification can be done automatically, the information for the tags of the rest of the collection will come out of the artificial neural network. Genus-, section- and species-tags were added to the pictures of the slipper orchids on the Flickr website prior to this project.

Adding tags to the pictures on Flickr was automated, as there are hundreds of pictures and multiple tags per picture. This was done by a Python script that makes use of the FlickrAPI [5, 6]. This API makes it possible to connect to a Flickr account from a Python script and make adjustments to data held by that account. For example: it is possible to upload or download pictures from the connected account, but also to add or remove tags of pictures.

Once the pictures on Flickr were tagged, the pictures with relevant tags (i.e. that identify the taxonomic group of interest) were automatically downloaded to become the training dataset and the metadata of these images were stored in a relational SQLite database [7]. There was a script available that did this for the orchid pictures, but it needed to be adjusted to work for butterflies as well.

### Image pre-processing and feature extraction

The slipper orchid photographs used previously had been taken in nature, so in the original OrchID framework the background of the orchid pictures was removed and only the flower of interest was retained, centred, and with any axial tilt corrected (this procedure is referred to as “image segmentation”). However, as all butterflies had been photographed in the same standard orientation at the same distance and on a fixed background, this was less of an issue than with the orchids, which for example had other plants in the background. This meant that the butterfly images could simply be cropped, but if the program is used for images without such standard orientation, segmentation needs to be done in addition to cropping. However, such segmentation was not part of this project.

When the correct part of the image, the Region Of Interest (ROI), was selected, features needed to be extracted. In the original framework, only the mean blue, green and red color intensities for a predefined number of rows and columns in the image were used for analysis, but for the butterflies a new type of features (Speeded Up Robust Features, or SURF) was added to the system. In SURF, features are algorithmically detected characteristics of the image, like corners and edges of stripes and spots. When a SURF feature is extracted from an image, two values become available: a key point and a descriptor. A key point is the coordinate set of the feature in the image. A descriptor is a vector of 128 elements to describe the feature and its surrounding area. For detailed information on how the descriptor is created, see *Appendix A* [8]. The key point itself cannot be used to compare images, because any rotation interferes with homology detection, but the descriptor can. The descriptors of the features were therefore used for image analysis. Upon feature extraction, the image was therefore represented by an array with the descriptors of every feature in the image.

Background removal and feature extraction were done in the original framework with OpenCV 2.4.x (www.opencv.org) and NumPy (www.numpy.org). Open Source Computer Vision (OpenCV) is a library of programming functions made for real time computer vision. It is written in C++ but has multiple interfaces to use it, among others in Python. It supports Windows, Linux, Android, iOS and Mac OS and has over 2500 available algorithms [9, 10]. The NumPy package contains the N-dimensional NumPy array and a set of mathematical functions. This package can be used for efficient array computations and manipulations, like calculations and comparisons of lists with thousands of descriptors of 128 elements each [11].

### Visual assessment of image feature performance

To visually assess species specificity of the features, Principal Component Analysis (PCA) was performed on the data. The code to do this was developed and executed by Wim van Tongeren, a Biology MSc student at Leiden University.

### Artificial neural networks (ANNs) were used to classify the new images

Artificial neural networks (ANNs) were used to classify the new images. An ANN is a machine learning algorithm that mimics the brain: input signals enter at input nodes, they are weighted according to their importance along nodes in one or more ‘hidden layers’ and combined together to create output signals at the output nodes [12]. In the ‘hidden layers’, various types of signal processing and topological rearrangements are applied to approximate the expected output signal: the classification. In this project the input signals were image features, and the networks were made using the Fast Artificial Neural Network Library (FANN) [13]. FANN is “a free open source neural network library, which implements multilayer neural networks for both fully connected and sparsely connected networks” [13].

The code for the new feature extraction methods needed to be implemented in the OrchID ANN-training framework, and some adjustments needed to be made to make the system work for both orchid and butterfly datasets. For example: the orchids were classified on ‘genus’, ‘section’ and ‘species’, but butterflies were only classified on ‘genus’ and ‘species’, because ‘section’ is a concept only in botanical taxonomy. A part of the data was used to train ANNs. The accuracy of a created ANN was validated by performing a stratified *k-*fold cross validation, like in the original framework. The validation of the ANN was done by Feia Matthijssen, a Biology MSc student at the University of Wageningen.

### Version control

Git is a version control system designed by Linus Torvalds, the creator of the Linux operating system [14]. GitHub is a website that makes use of Git to work in groups on software development. On the website (www.github.com), source code, data and documents are stored in repositories. Each repository ideally has its own topic with corresponding documents, data and source code. GitHub works with accounts, and every account can have multiple repositories. Depending on the rights of a repository, people can change files in the repositories of another person’s account, so you can work together on a project [15].

The output files and scripts of this project were stored in repositories of the Naturalis organization on GitHub. The (interim-) output files were stored in the repository called ‘nbclassify-data’ (https://github.com/naturalis/nbclassify-data), the scripts and modules in the ‘nbclassify’ repository (https://github.com/naturalis/nbclassify). The repository ‘imgpheno’ (https://github.com/naturalis/imgpheno) is a package that makes use of the scripts in ‘nbclassify’. These repositories are grouped together with other image recognition projects of the Naturalis Biodiversity Center in the ‘img-classify-all’ repository (https://github.com/naturalis/img-classify-all).

### Goal of the project

Add image features to the existing framework ‘OrchID’ to identify specimens of the Javanese butterfly collection and to make the recognition of slipper orchids more specific.

## Materials and methods

### Project workflow overview

*Figure 1* shows a flowchart of the steps taken during this project for the butterfly part. The taxonomy of the butterflies and images on Flickr were available at the start of the project, but more data became available during the project, so the workflow up to the clustering was repeated multiple times. Development was done in Python 2.7.11 and R version 3.2.3. All outcomes (code and result files) were placed under version control in repositories of the Naturalis GitHub organization. All orchid pictures were already tagged on the Flickr website prior to this project. Therefore the first part of this flowchart could be skipped for the orchid part of the project. From downloading onward, the workflow was identical for the orchids.

**Figure 1.**
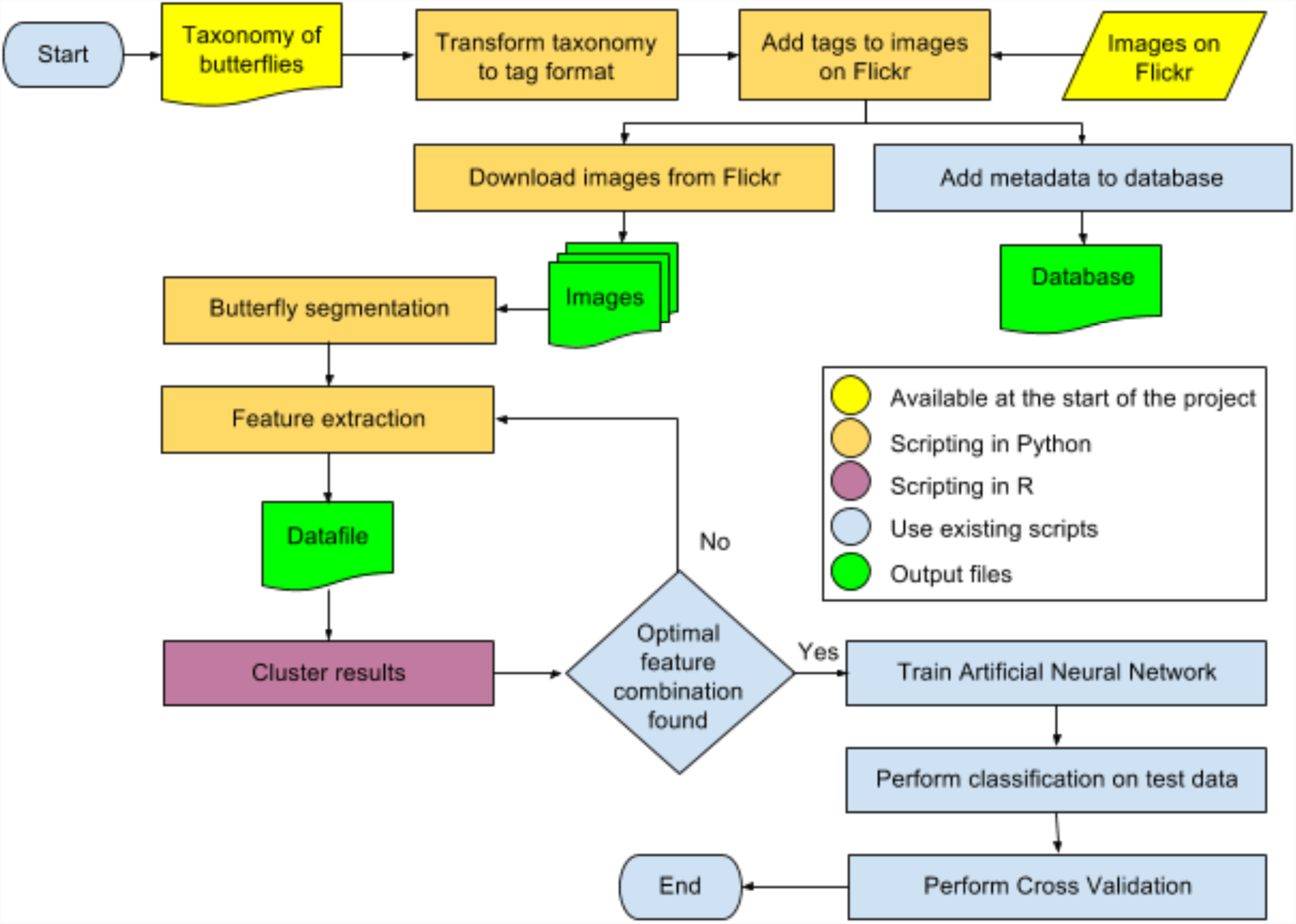
Flowchart of steps that were taken in this project. This flowchart applies to the butterfly analyses of this project. From the download-step onward, it applies to the orchid analyses as well.

### Working environment

To make use of the existing modules and scripts that were stored on GitHub, the imgpheno and nbclassify repositories needed to be installed according to the commands in the README.rst files in these repositories. For the GitHub connection, Git version 2.7.4 was used. The installation commands were executed from the Git shell.

*Figure 2* gives an overview of the contribution of these repositories, among other modules, to the project. The OrchID web application was not developed further in this project.

**Figure 2.**
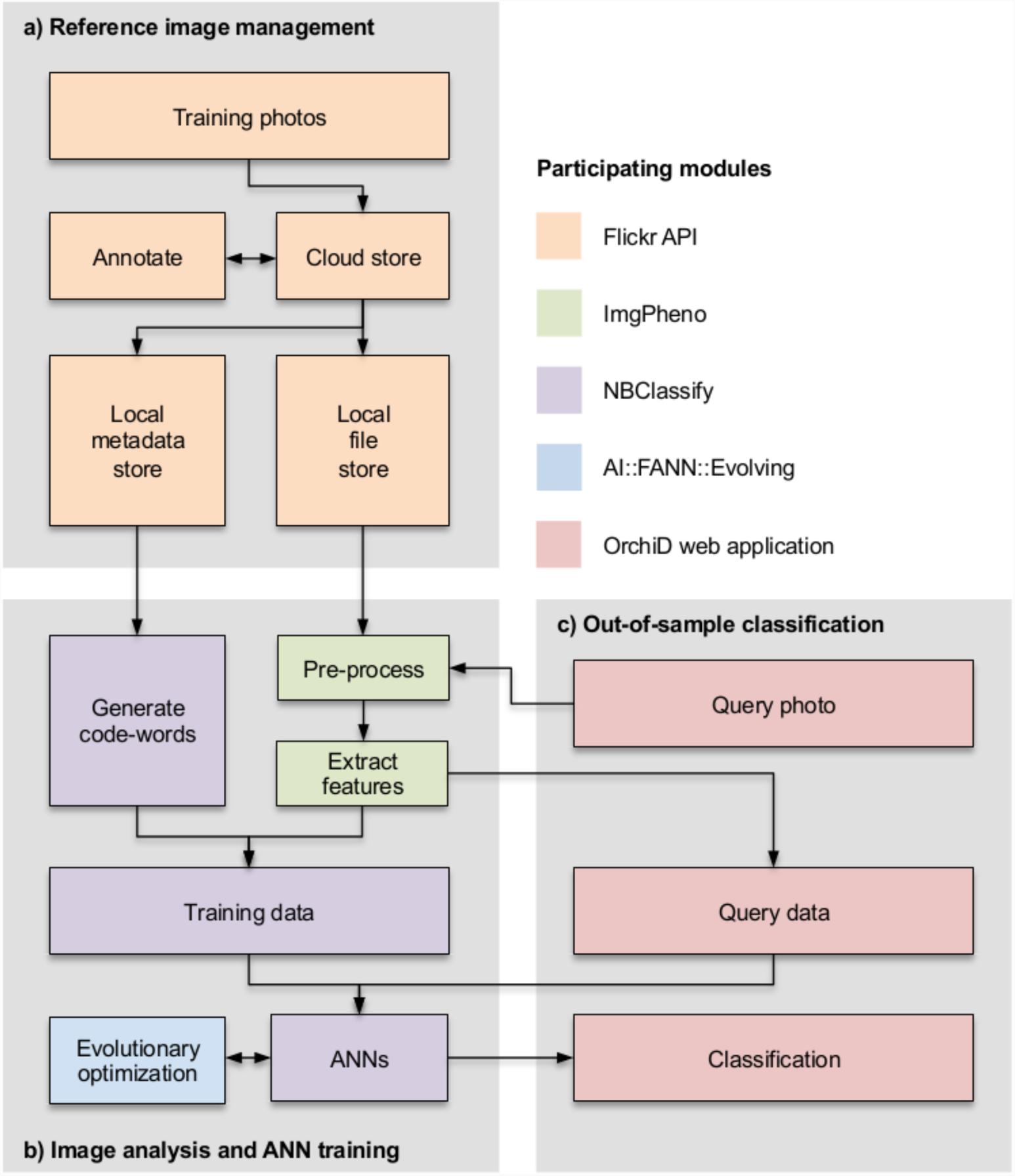
Overview of the participating modules in this project. The OrchID web application was not used in this project.

All of the code involved in this project was adapted to work on both Windows and Linux systems. At first, the whole process was done on a local Windows machine, but later on a Linux-based cloud environment became available for the project. The cloud had four cores, a storage space of 1TB, 64GB RAM and 2.60GHz CPU per core. A connection to the cloud was made on the Windows machine by using PuTTY: “a free SSH and Telnet client for Windows and Unix” [16]. PuTTY was already installed on the local computer prior to this project, but is available from the PuTTY Download page [16]. The instructions for the connection and used parameters are described in *Appendix B*. The session was saved with the name ‘Butterflies’, so this could easily be loaded every time a connection to the cloud was desired. Although the name suggests otherwise, the same connection was used for the orchid part of this project.

Files were transferred from a local computer to the cloud and the other way around using the ‘pscp’ client, available at the same website as the PuTTY client [16]. Because the session was saved as ‘Butterflies’ in PuTTY, this name could also be used to make the connection with the pscp command. The pscp commands were all performed by a command line interface in Windows. See *Appendix C* for pscp usage instructions.

### Image data and metadata management

The butterflies of the Javanese butterfly collection were stored in papillottes in drawers. Each specimen was taken out of the papillotte and photographed in standardized position on a paper with a ruler on it. This was done by staff of the Collection department of Naturalis Biodiversity Center. In each picture, on the top right there was a card with information known about the specimen, such as the place where it was collected, a collection registration number and a QR-code to the collection registration system (CRS) of Naturalis Biodiversity Center. If there was any information about the specimen written on the papillotte, such as a date, this papillotte was put in the lower right corner. The specimen was placed on the left side of the paper facing the right. It was placed in the position it was stored all those years, so (except for a few specimen) with folded wings. This meant that the underside of the wing (ventral side) was visible in the picture. *Figure 3* shows such a picture.

**Figure 3.**
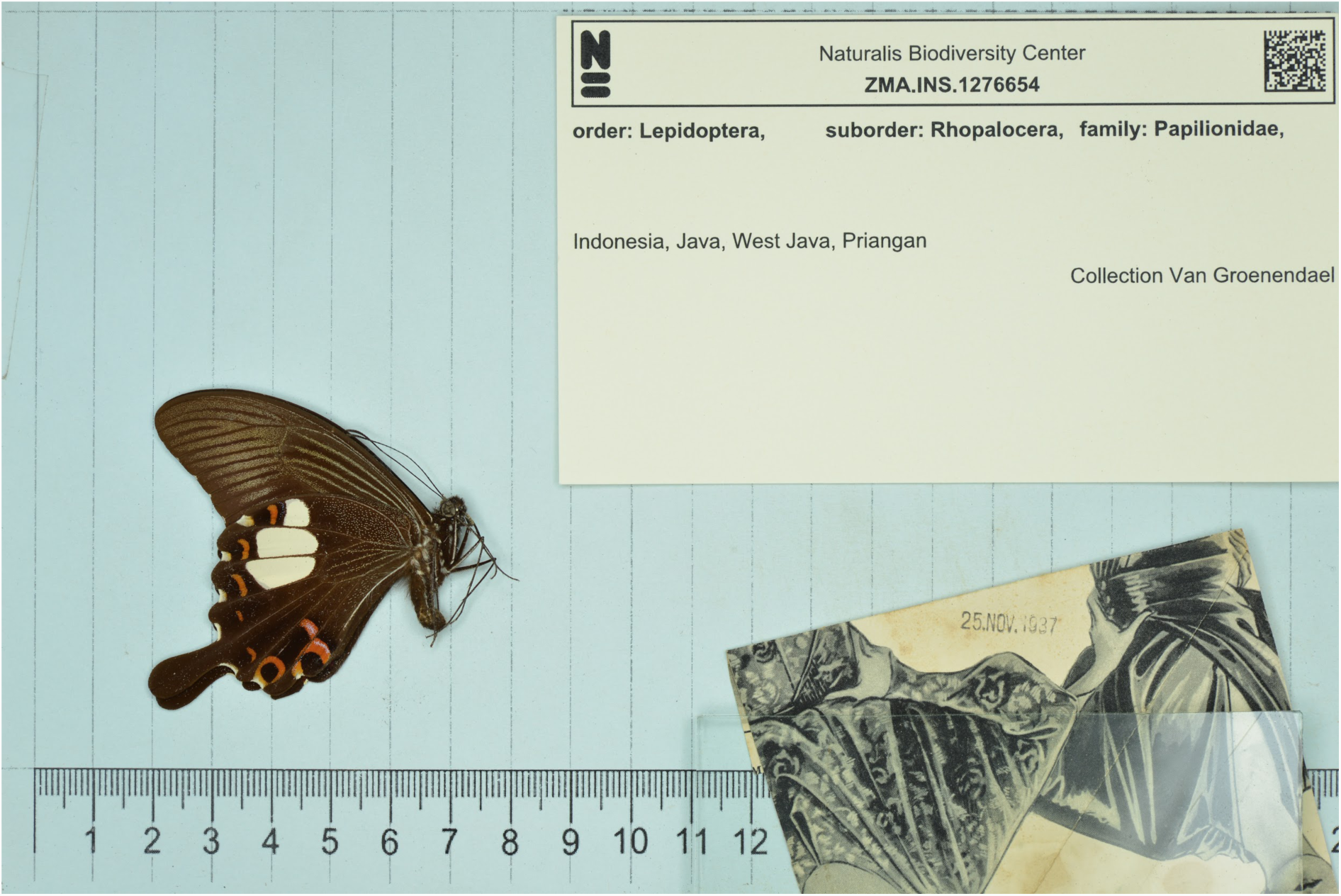
Standardized picture of a specimen of the Javanese Butterfly collection. In the right upper corner there is a card with metadata of this specimen. In the lower right corner there is the papillotte with a date on it. On the left side there is a specimen with folded wings, facing the right. All items were placed on a paper with a ruler in the bottom.

Butterfly taxonomist Jan Moonen classified almost 1800 pictures out of the Javanese butterfly collection, divided over three datasets. The pictures he had classified were uploaded to an account associated with the project on the Flickr website (www.flickr.com), using the upload option on the website. The information provided by Jan Moonen for every picture was listed in a Microsoft Excel spreadsheet. Sometimes only the ranks family, genus and species were given, without the ranks in between, so they were looked up in the Insects Catalog on the website of the International Entomological Community [17] and added to the sheet. Some of the column headers and remarks in the spreadsheet were in Dutch, so they were translated into English. All headers were changed to lowercase and the male and female symbols used (♂ and ♀), were changed into M and F respectively.

In the three identification files, the name of the identifier had a mixed format. The format of the first file was chosen to use in every other file: “Moonen, J.J.M.”. All other formats were changed into this one. The identification date had a Dutch format: DD-MM-YYYY. This was changed into the ISO 8601 standard format: YYYY-MM-DD [18]. The file was exported as a CSV file. Genus, subgenus, species and subspecies were in *Italic*, but this was undone by the conversion to plain text CSV format, which also required changing the Unicode symbols for male and female to simple letters. The remarks of the third identification file indicated that there were some damaged specimens in this dataset. After investigation it was decided to leave eight specimens out of this dataset, so they were removed from the Flickr website.

A Python script was made to change the spreadsheet information into tag format compatible with Flickr (example: ‘genus:Papilio’) and to automatically add these tags to the pictures on the website. The script made use of the FlickrAPI [5] to annotate the pictures on the Flickr account. The FlickrAPI needed a Python version like > 2.7.9. To connect to the Flickr website, different methods are available. The method used in this script was ‘authenticate via browser’, which needs FlickrAPI version 2.1.2 (see *Appendix D* for download and installation instructions). The first time a script with ‘authenticate via browser’ is run, a webpage opens and a button needs to be clicked to verify the connection. Every subsequent time, authentication is done automatically.

For all scripts in this project, the available arguments can be viewed with the -h option, like:

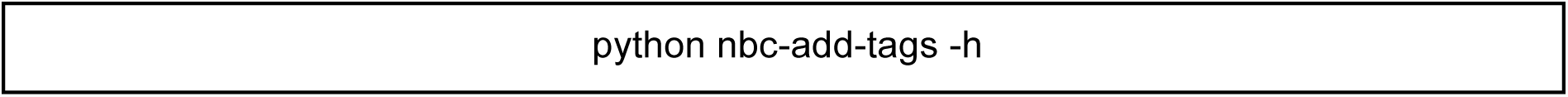

The script ‘nbc-add-tags’ was used to add tags to the pictures. The arguments that were used to call this script are described in *Appendix E*. The script was run for all three butterfly datasets.

The orchid pictures came from different collectors and orchid specialists, who also provided the names of the flowers in the pictures. The pictures were taken in nature, so without a standardized setup. Prior to this project these pictures were segmented, so the flower was in the centre of the picture and the background was as much as possible removed. *Figure 4* shows such a picture. These segmented pictures were already uploaded to the Flickr website and tagged with a family-, genus-, section- and species tag.

**Figure 4.**
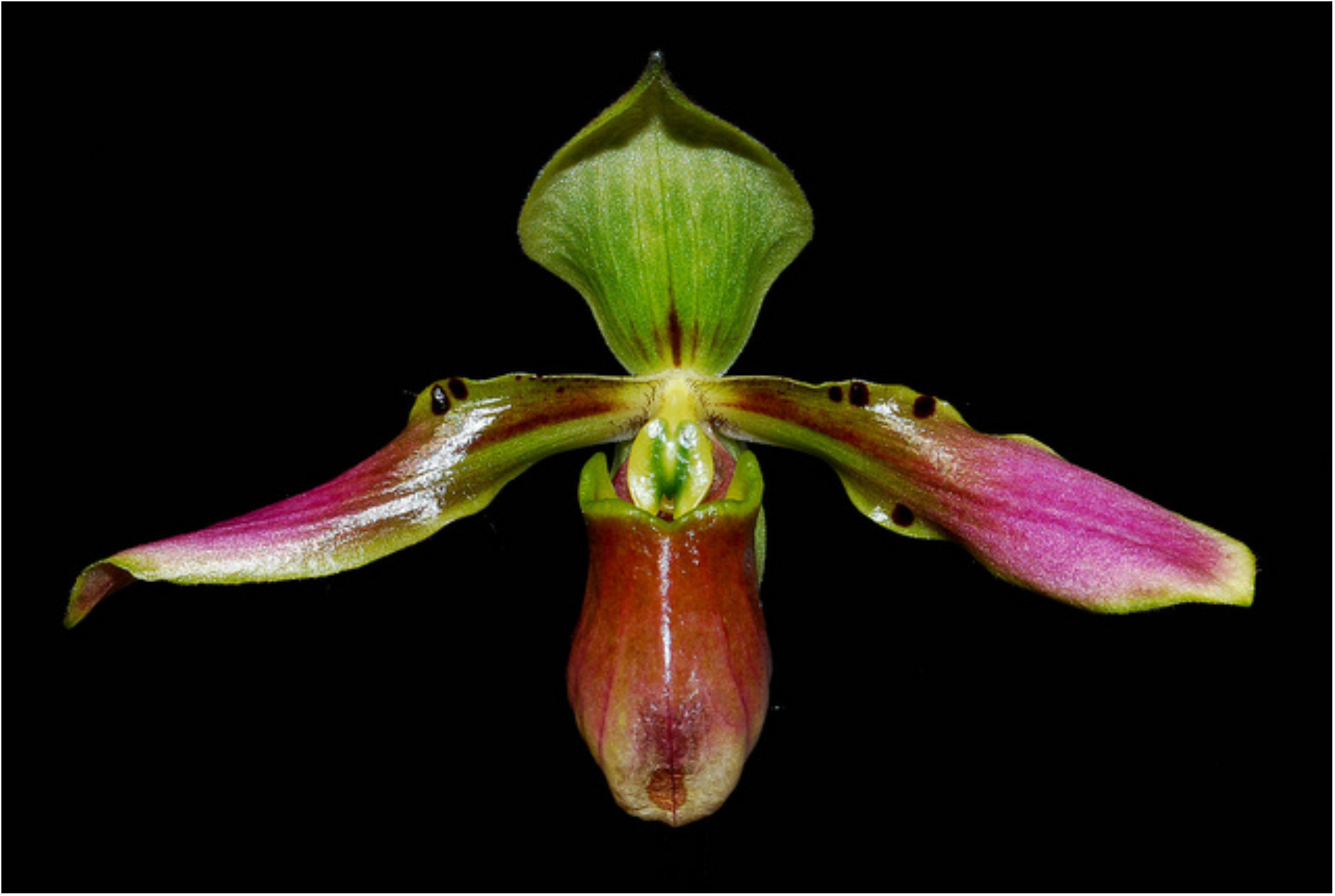
Picture of a specimen of the slipper orchid dataset. The flower was placed in the center and the background was automatically removed prior to this project.

Relevant pictures were downloaded from the website. A butterfly picture was considered relevant, when the genus and species tag were set, no matter what other tags were set as well. For the orchids the section tag also had to be set. To download the relevant pictures from the website and save the metadata in a database, the existing script “nbc-harvest-images” was adjusted and used.

Various arguments were added to the argument parser, to make it possible to give some extra arguments to the nbc-harvest-images script instead of using hardcoded variables. The script makes use of the tags on the Flickr website, to decide whether or not to download a picture. There was already an option to give a complete tag as argument to the script and only select those pictures that contain the complete tag. For example, only download the pictures that belong to the genus *Papilio*. Now the option to select pictures based on only the key-part of a tag was added. So, for example, only download pictures that have a ‘genus’ tag, irrespective of what the genus is. This was necessary because otherwise the script should be called for every genus separately. There was also an option added to decide if the pictures would be saved in a <genus><species> directory hierarchy or to use all possible taxonomic ranks for the directory hierarchy.

This script makes use of the FlickrAPI like the nbc-add-tags script, but was created with another authentication method: ‘authenticate via token’. This older method needs FlickrAPI version 1.4.5, so this was installed over the FlickrAPI 2.1.2. In the future this needs to be changed, so that both scripts will use FlickrAPI version 2.1.2. Download and installation instructions are given in *Appendix D*.

The nbc-harvest-images script connects to the Flickr website and harvests the pictures that meet the requirements set in the arguments of the command to run the script (see *Appendix F* for used arguments). Once a picture is harvested, all its tags, photo ID, photo title and photo md5sum (digital fingerprint [19]) are stored in a relational database. If there is no existing database available, a database is automatically created by the script “db.py” that makes use of the sqlalchemy library [20] to create and communicate with a relational (SQLite) database. The sqlalchemy library is automatically installed when the nbclassify repository was installed. There has been a minor change to the ‘db.py’ script as well. A check was added to verify the type of a file that was about to be entered into the database. If it was not of type np.ndarray, the file was not a picture and therefore must not be entered.

Both butterflies and orchids had a genus and species tag, so when these tags are the only requirements, they would both be downloaded instead of only the butterflies. The id_date tag was provided for all the butterflies, but not for the orchids. Therefore it was used with genus and species as compulsory tag to distinguish the butterfly pictures from the orchid pictures on the Flickr website. The butterflies do not have a section tag, so only orchids were downloaded when this tag was required.

The script was run for all three butterfly datasets and the orchid dataset. Each separate butterfly dataset was downloaded before the next dataset was uploaded. The newest pictures on the site are always on the first page, so the datasets could be downloaded from the first page onward. Now all datasets are uploaded to the website, a starting page number and perhaps another number of pictures per page have to be given to download a separate dataset again. The script was run a fifth time with only <genus><species> directory hierarchy for the three butterfly datasets at once. This was necessary because the scripts that make the artificial neural network, are designed to work with this hierarchy.

### Reference data selection

Once the third butterfly dataset from Jan Moonen had come in, inventory was taken to count the number of specimens of every species in the reference dataset. Some species had a morphological difference between sexes, so each of those sexes was treated as a different species. It turned out that there was a great difference in number of specimens per species (between 1 and 267), so a selection of the data was made to create a reference set with a more evenly distributed species count. Species with fewer than 5 specimens were left out of the selection. A maximum of 50 was taken for the other species. The pictures were randomly chosen but visually inspected to make sure the butterflies in this selection were of good quality and damaged specimens or specimens that were folded in the wrong way were excluded from the dataset.

The selected dataset was used to train the artificial neural network (ANN). A stratified *k*-fold cross validation was performed to test the created network. After validation of this network it was decided to leave species with less than 15 specimens out of the selection as well. With this final dataset, a new ANN was created for the classification of the rest of the Javanese butterfly collection. Lists with the pictures used in the first and definitive selection were stored in the nbclassify-data repository in the archive-butterfly directory.

For the orchids a similar inventory was taken. This dataset was limited in the number of specimens per species. Here also the minimum was set to 5, the maximum to 50, but the species with the most pictures had only 36 specimens, so this maximum was never reached. With this dataset the orchid ANN was created and validated.

### Image feature extraction

Features were extracted from every butterfly and stored in a spreadsheet; thereafter attempts were made to cluster specimens by their extracted features to assess whether the applied method gave a species-specific output. Feature extraction and clustering were done iteratively, in order to get the best available method to identify the species in the picture. This method was then also applied to the orchid pictures to determine if this method was better than the original method for taxonomic identification of slipper orchids.

To perform image feature extraction on the butterflies, a fixed Region Of Interest (ROI) was determined in such a way that the same ROI could be applied to all standardized butterfly pictures. This was done manually by making a subset of the first dataset with pictures of the largest specimens and the specimens that were placed more to the sides than others. The region that contained all specimens was determined by cropping the images until all specimens were included and as much background as possible was excluded. Every new dataset was examined on extreme positions of the specimens and, if necessary, the ROI was re-determined. With the first dataset, a ROI was determined in pixel units, but in the second dataset some pictures had a different resolution, so the pixel units could not be applied. The pixel units were then transformed into fractions of the whole picture size, so the ROI could be applied to all standardized pictures. The orchid pictures had already been segmented, so a ROI for these pictures was not necessary.

In the original OrchID framework, an image was divided into 50 horizontal and vertical bins and the mean blue-green-red (BGR) values were calculated. This method was also applied to the butterfly pictures. Although this method was used for the orchids, there was no script available that could automatically apply the method to the pictures. A Python script called “nbc_create_mean_bgr_file.py” was therefore created, that made use of the available methods in the imgpheno package. The script also used the database created with the nbc-harvest-images script, to create a tab separated spreadsheet with the photo ID, photo title and all horizontal and vertical normalized mean blue, green and red intensity values per bin. This script was run only for the first two datasets, to verify if this method would work for the butterflies. *Appendix G* describes the used arguments. Since this method was used for the orchids in the original framework, this was not repeated in this project.

Another method was also applied to extract features from the butterfly and orchid images: the Speeded-Up Robust Features (SURF) algorithm of OpenCV [8, 21]. This algorithm detects changes in pixel intensity to locate features (e.g. corners and edges of stripes and (eye)spots) of an image [21]. OpenCV version 2.4.6 was installed, because it included this SURF algorithm for free. Newer OpenCV versions for Windows also include this algorithm for free, but for Linux it became patented and now needs to be paid for [22]. See *Appendix D* for download and installation instructions of OpenCV version 2.4.6.

The Python script ‘nbc-feature-extraction.py’ was created to extract features using the SURF algorithm. In this script all files of the (subdirectories of the) base directory were read and checked to be an image. If so, the image was cropped to the ROI if necessary and converted to gray scale, because the SURF algorithm needs an image in grayscale to detect changes in intensity. The butterfly images were truncated to fade the lines of the ruler in the background, otherwise the lines would be seen as features as well. To do so, the OpenCV method cv2.threshold() was used with cv2.THRESH_TRUNC as threshold style [23]. A threshold value of 170 was used: all pixels with an intensity value of 170 or above were given a new pixel intensity value of 170, so the brightest parts of the image would turn a little darker. In that way the differences between the background color and the lines of the ruler were too small to detect by the SURF algorithm. The orchid images were not truncated.

The descriptors of the features that were detected by the script were stored in a dictionary, with the filename as key and the descriptors as value. When all pictures were processed, the dictionary was saved to a file with the dump method of cPickle [24]. In *Appendix H* the used arguments for nbc-feature-extraction.py are described. This script was run for the first two butterfly datasets and the orchid dataset. It was not run for the third butterfly dataset, because this script was only used to test the species specificity and the results of the first two datasets were clear enough.

Each image had a different number of features, even if it was the same species, so it was impossible to compare the descriptor arrays without a form of data transformation and reduction. To make it possible to compare the features, the Bag-Of-Words (BOW)-method was applied to the file with the dictionary of descriptors. This method is commonly used in document classification, but more and more in computer vision as well [25-27]. For a detailed explanation with examples of the steps of the Bag-Of-Words method, see *Appendix I.*

The Python code ‘nbc-bag-of-words.py’ was created to apply this method to the files with descriptors. The code in the GitHub repository ‘Minimal-Bag-of-Visual-Words-Image-Classifier’ on the account of Ludwig Schmidt-Hackenberg (accountname ‘shackenberg’) [28], was used as an example to create the ‘nbc-bag-of-words.py’ script. In this script, a *k*-means clustering was performed on all 128 elements of all the feature descriptors in the dictionary file (so in 128 dimensions). The codebook was created with the centroid positions of every cluster, and every centroid was described by an array of 128 elements.

For every image it could then be determined to what cluster each feature belonged by comparing them to the codebook, resulting in a vector per image with the centroid index numbers in the codebook for every feature. These vectors have the length of the number of features in that image, so a histogram was made of every vector compared to all the centroids in the codebook. This led to word-histograms: a vector per image of length equal to the number of centroids in the codebook. The counts in this histogram were corrected for the number of features in the image with the fraction of times every centroid was represented in that image as a result.

The word-histograms of all images have the same length: the length of the codebook, hence the number of *k*. New images that were not in the dataset that created the codebook could also be compared to the existing codebook, so the word-histogram of a new image could be compared to already known images [29]. The codebook was saved with the dump method of cPickle. The word-histograms were saved in a tab-separated file, with the photo ID, photo title and the fraction each centroid appeared in the image. The database that was created in the nbc-harvest-images script was used to get the photo ID and photo title.

The output of the nbc-feature-extraction.py script was used as input for the nbc-bag-of-words.py script, so it was run for the first two butterfly datasets and the orchid dataset, like the nbc-feature-extraction.py script. For the first and second dataset, the script was run with a *k* value (argument --clusters) of 150, 500 and the square root of the total number of features, in order to get the optimal one, i.e. the lowest number that was still species specific. It was decided to use *k* = 150 as optimal one. The orchids had too little features to use *k* = 150, so *k* = 125 was used. For the other used arguments, see *Appendix J*.

### Visual inspection of image feature performance

In order to assess visually the performance of different image features in terms of their ability to cluster specimens correctly, PCA was performed. The R code for this clustering was created and executed by Wim van Tongeren, a MSc Biology student. In this code, various statistical analyses were done to determine the need for normalization and data transformation in order to perform a Principal Component Analysis (PCA) on the data. The results of the PCA were visualized by plotting principal components one and two, one and three and two and three, each on a two-dimensional (2D) surface. These three plots were made to inspect the 3D position of the data points. In a 2D plot, data points may look like they are grouped together, but when examined from another angle they might be in separate groups behind each other. The code was saved as .Rmd file in the nbclassify-data repository on GitHub, in the archive-butterflies directory (https://github.com/naturalis/nbclassify-data/blob/master/archive-butterflies/butterfly150.Rmd).

All butterfly plots were made with both genus and species as label and were visually inspected for, respectively, the genus- and species specificity of the applied method that created the data. For the orchids, also the section-specificity was inspected. In order to be genus-, section- or species specific, ideally all specimens of one genus, section or species group together in the PCA plot. When a specific data point needed more attention, for example to clarify why it was not grouped with other members of the group it belonged to, the photo ID or photo title were used as a label in the plot, so the original image could be traced.

The spreadsheets that came out of the script ‘nbc-create-mean-bgr-file.py’ for the first two butterfly datasets were each used for a PCA. The data was plotted to determine if the method that was used in the original OrchID framework for the slipper orchids, would also give genus- and species specific results for the butterflies and could therefore successfully be applied to both slipper orchids and Javanese butterflies.

The word-histograms from the SURF-BOW method were plotted to determine if this method was in any way genus-, (section-), and species specific. The word-histograms of the butterflies were also used to determine what the best *k* value would be for the *k-*means clustering in the BOW-part. The used *k*-values were 150, 500 and the square root of the total number of features. It was decided to use the *k*-value with smallest value that was still genus- and species specific.

### Artificial neural network training

Once the image feature extraction methods had been developed and applied to the image sets, artificial neural networks (ANNs) needed to be trained on these data. The scripts to train, validate and use ANNs were created for the original OrchID framework and could be reused for the butterfly part of the project. These modules were made to do all the necessary steps of the image analysis from the point where the dataset was available in a directory, so including the pre-processing steps like background removal and segmenting the image, but the image harvesting from Flickr was excluded. There were 11 modules in the nbclassify repository that collaborated with each other and with parts of the imgpheno package in order to make a functional ANN, based on the mean BGR values of an image.

To be able to create an ANN based on the SURF-BOW method, the code to extract image features and cluster them according to this method was added to the existing ANN modules. Therefore, the code of the scripts ‘nbc-feature-extraction.py’ and ‘nbc-bag-of-words.py’ was implemented in the scripts for the ANN. The scripts ‘nbc-feature-extraction.py’ and ‘nbc-bag-of-words.py’ can still be used to create interim output files, which are only created with the validation scripts for each species separately, but it is not necessary to use them for the ANN. A config.yml file was made to set all the parameters for the butterfly- or orchid identification. In this config.yml file was described what method(s) should be used for the feature extraction. First, only the SURF-BOW method was used, but later on both mean BGR values and SURF-BOW methods were applied to the pictures to check if the network would be better with both methods combined instead of only one.

### Artificial neural network cross-validation

ANNs may appear to perform well on a given combination of training data and out-of-sample date solely due to overfitting. Hence, the general performance of an ANN training and classification approach needs to be validated across data sets. Such validation of ANNs was done by Feia Matthijssen, a Biology MSc student of the University of Wageningen. According to her findings (https://github.com/naturalis/nbclassify-data/tree/master/archive-butterflies/nbc-validate-results) adjustments to the scripts were made. The validation included the iterative partitioning of data into training (a fraction of *k*-1/*k*) and testing (1/*k*), then training the ANN, and quantifying the performance on the test partition, i.e. stratified *k*-fold cross validation. These steps were all built in the original scripts and could be called by giving different arguments to the argument parser of the script ‘nbc-trainer’. Because of memory constraints and duration of the Bag-Of-Words-part, this script was always run on the cloud. The numbers of *k* used for the stratified *k*-fold cross validation were 2, 5 and 10 for the butterflies and 5 and 10 for the orchids.

The system was first tested by making use of a very small subset of 15 butterfly pictures. Afterwards the network was validated with the first two butterfly datasets combined, but after three days the program with the validation task was still running without any progress, due to the high number of features that had to be clustered into a high number of clusters for the Bag-Of-Words method, so it was killed. Then the selected butterfly dataset, with only species that had between five and 50 specimens, was used for the validation to make sure there was a strong dataset to train the network with and also to lower the number of items in the dataset in order to speed up the validation process. The selected butterfly dataset was used for the validation of a network based on only the SURF-BOW features and to validate both methods combined. According to the results of these validations, the definitive butterfly dataset was made and the final network was created to use for the classification.

The selected orchid dataset, with only species that had between five and 50 specimens, was used to validate the network that was created with the SURF-BOW feature extraction method and for both methods combined. These results were compared to the results from the original OrchID framework with only the mean BGR values to find the best method for the identification of the orchids in this dataset.

### Version control

To make sure the latest version of the GitHub repository was used, a synchronization with the website was done before a script was changed by pulling the information from the website. This was done for the nbclassify, imgpheno and nbclassify-data repositories. All latest versions of the documents in the repositories were then locally available. After changes were made to a file, this file needed to be pushed onto the GitHub website. The pulling and pushing was done on a commandline. See *Appendix C* for the general commands for these actions. Documents could also be uploaded by the ‘upload files’ button on the website.

When changes were made to the nbclassify or imgpheno repository, the installation of the repository had to be upgraded. The command to do this is also described in *Appendix C*.

## Results

### Image data and metadata collection

The original butterfly identification files Jan Moonen provided are stored in the nbclassify-data repository in the directory archive-butterflies/Identification_files. The following changes were made to these files: header to lowercase, no *Italic*, male/female symbols changed to characters, format of the name of the identifier changed, identification date in YYYY-MM-DD format and all Dutch translated into English. *Table 1* displays a part of an identification table after the changes were made. These new tables were saved as .CSV files and are stored in the same directory as the original files.

**Table 1.** Part of an identification file after editing. Content is the same as the ongmal table, but now all headers are lowercase, there are no Dutch nor Italic items, the sex symbols are changed into characters and the format of identifier-and id date columns is changed

| reg_number | family | subfamily | tribus | genus | subgenus | species | subspecies | sex | identifier | id_date | remarks |
| --- | --- | --- | --- | --- | --- | --- | --- | --- | --- | --- | --- |
| 1277414 | Papilionidae | Papilioninae | Papilionini | Papilio | Achillides | paris | gedeensis | M | Moonen, J.J.M. | 2016-04-15 |  |
| 1277415 | Papilionidae | Papilioninae | Papilionini | Papilio | Achillides | paris | gedeensis | M | Moonen, J.J.M. | 2016-05-08 |  |
| 1277416 | Papilionidae | Papilioninae | Papilionini | Papilio | Menelaides | helenus | enganius | M | Moonen, J.J.M. | 2016-05-08 |  |
| 1277417 | Papilionidae | Papilioninae | Papilionini | Papilio | Achillides | paris | gedeensis | M | Moonen, J.J.M. | 2016-05-08 |  |
| 1277418 | Papilionidae | Papilioninae | Papilionini | Papilio | Menelaides | demolion | demolion | M | Moonen, J.J.M. | 2016-05-08 |  |

Originally, the first butterfly dataset contained 382 pictures, the second one 859 and the third one 612. The specimens that were deleted from dataset three and the reasons to delete them are listed in *Table 2*. After deletion, the third dataset contained 604 specimens.

**Table 2.** Deleted specimens from butterfly dataset three. The reason to delete the specimen is described

| Registration number | Reason |
| --- | --- |
| 1277820 | Damaged specimen |
| 1277823 | Damaged specimen |
| 1277890 | Damaged specimen |
| 1277918 | All loose parts |
| 1277930 | Damaged specimen |
| 1277960 | Damaged specimen |
| 1278012 | Folded in the wrong way |
| 1278013 | Damaged specimen |

The nbc-add-tags script ran three times. The first time it tagged 383 pictures, the second time 859 pictures and the third time 604. The nbc-harvest-images script ran also for each butterfly dataset. The first time it downloaded 373 pictures, the second time 839 and the third time 604. The pictures that were not downloaded, lacked a genus and/or species tag. For the orchid dataset it downloaded 1117 pictures. The fifth time it ran, the whole butterfly dataset was downloaded in a <genus><species> directory hierarchy. It took one hour and 59 minutes to download 1816 pictures. The taxonomy and number of specimens per species of the three butterfly datasets is listed in *Appendix K*. When both sexes of a species were in the dataset and they showed sexual dimorphism, they were treated as different species. Therefore the sex was added to the species name.

Some species had only one sex present in the dataset. *Atrophaneura nox* had only female specimens. *Graphium antiphates, G. bathycles, G. doson, G. eurypylus, G. evemon, G. macareus, Lamproptera curius, Papilio karna, P. paris, P. peranthus* and *Troides helena* had only male specimens. Those species could not be checked for sexual dimorphism. This may be done in the future to decide if they should be treated as different species as well.

In *Figure 5* the number of specimens per butterfly species of the three datasets combined is shown. In the selected butterfly dataset, all species with fewer than five specimens were left out (see the red cut-off line in *Figure 5)* and a maximum of 50 was used. Families *Pieridae* and *Riodinidae* had too few specimens per species to be selected. A definitive dataset was made with at least 15 specimens per species (the green line in *Figure 5* indicates this cut-off). The definitive butterfly dataset contained 617 pictures of 17 species in five genera.

**Figure 5.**
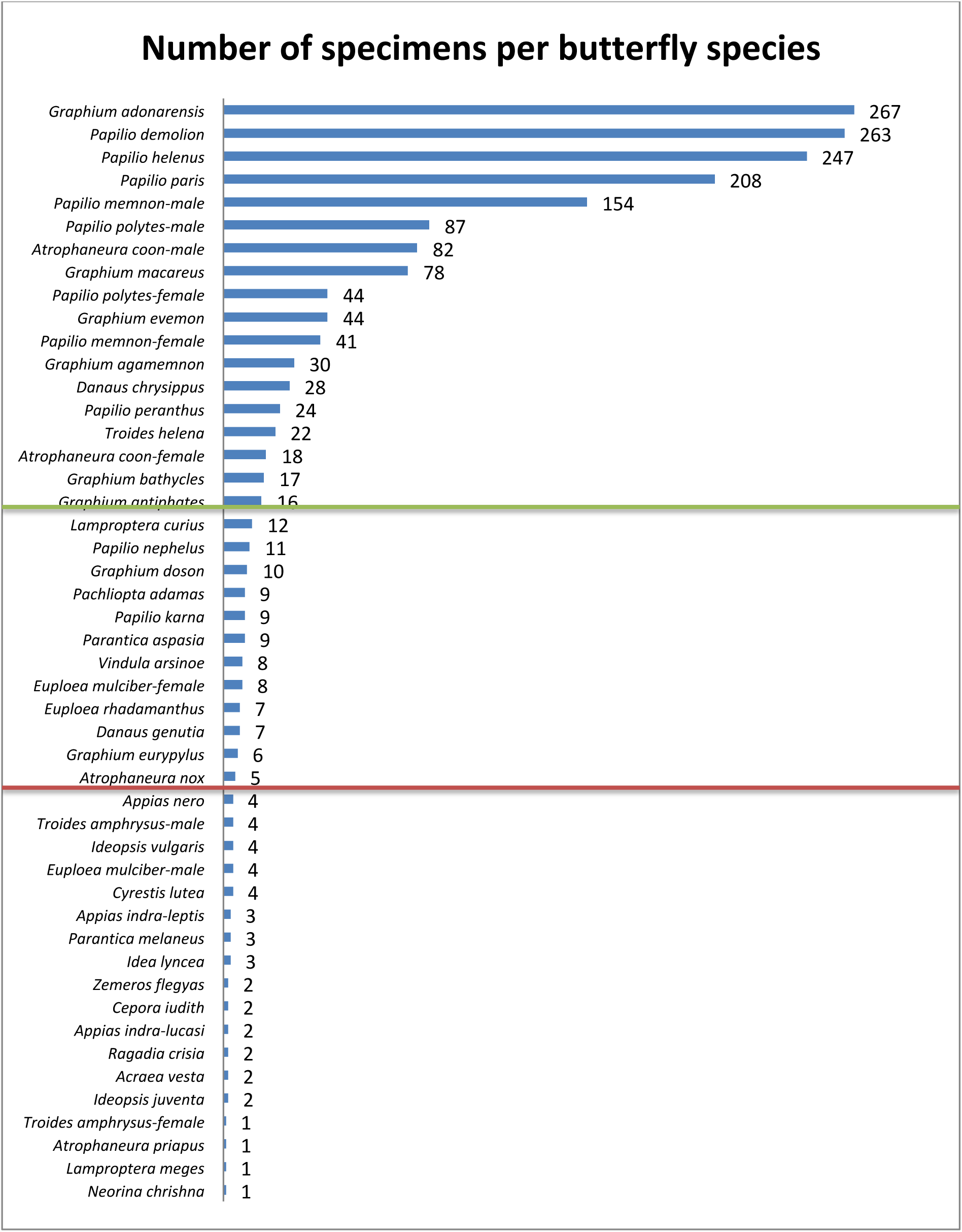
Number of specimens per butterfly species. The numbers are the total of the three datasets combined. The red line indicates the cut-off for the selected dataset (n=5), the green line indicates the cut-off for the definitive dataset (n=15).

*Figures 6-7* show the overview of specimens per species for the orchid dataset. There were five genera in this dataset. The species that had fewer than five specimens were left out of the selection and are located below the red cut-off line in *Figure 7*. The selection contains 1041 specimens of 84 species in 18 sections of four genera. The genus *Selenipedium* had only four specimens, so was left out of the selection. The green line in *Figure 6* shows where the cut-off would have been if a definitive orchid dataset was made with at least n=15, like with the butterflies.

**Figure 6.**
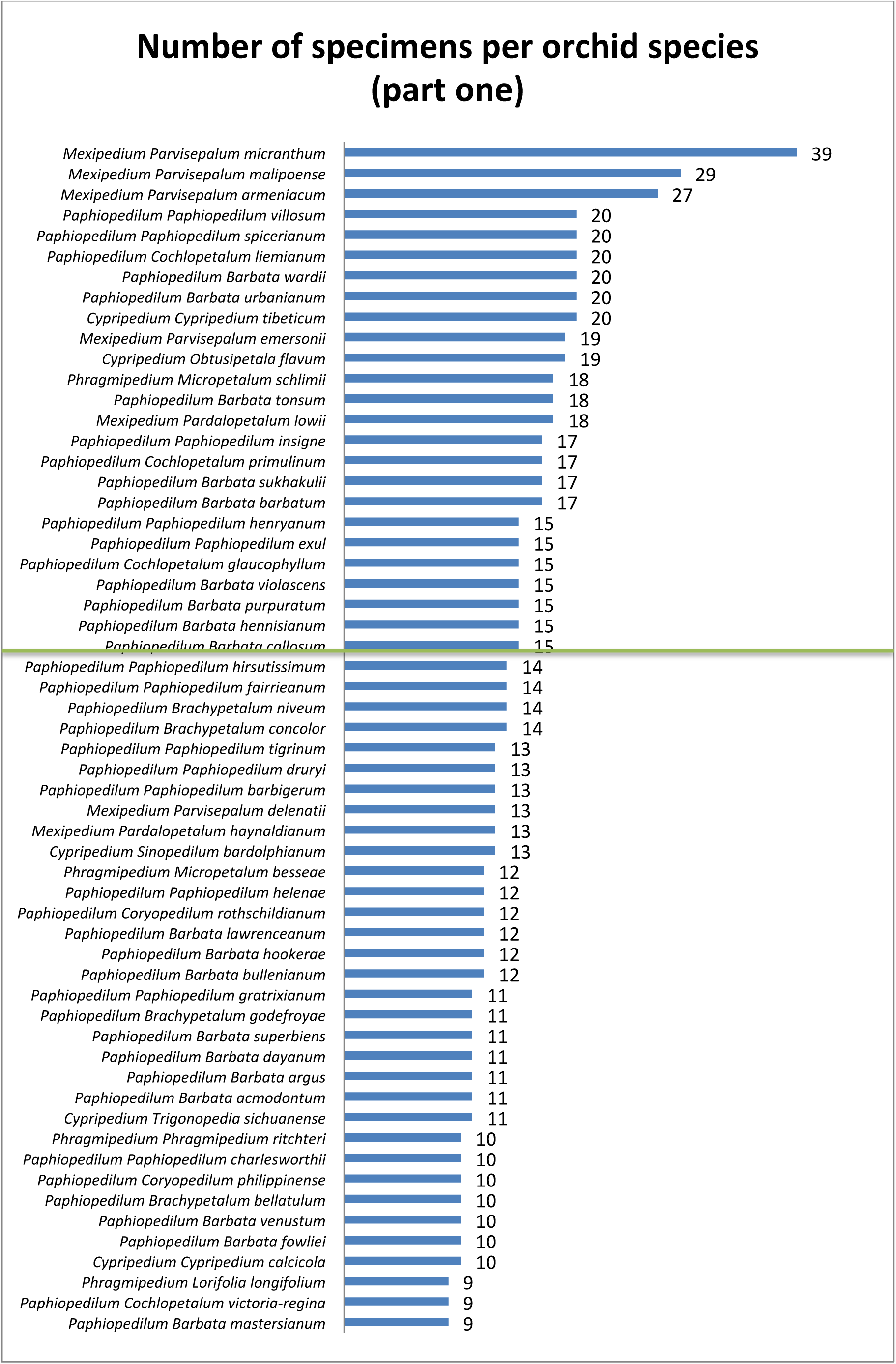
Number of specimens per orchid species part one. The green line indicates where the cut-off would have been for the definitive dataset (n=15).

**Figure 7.**
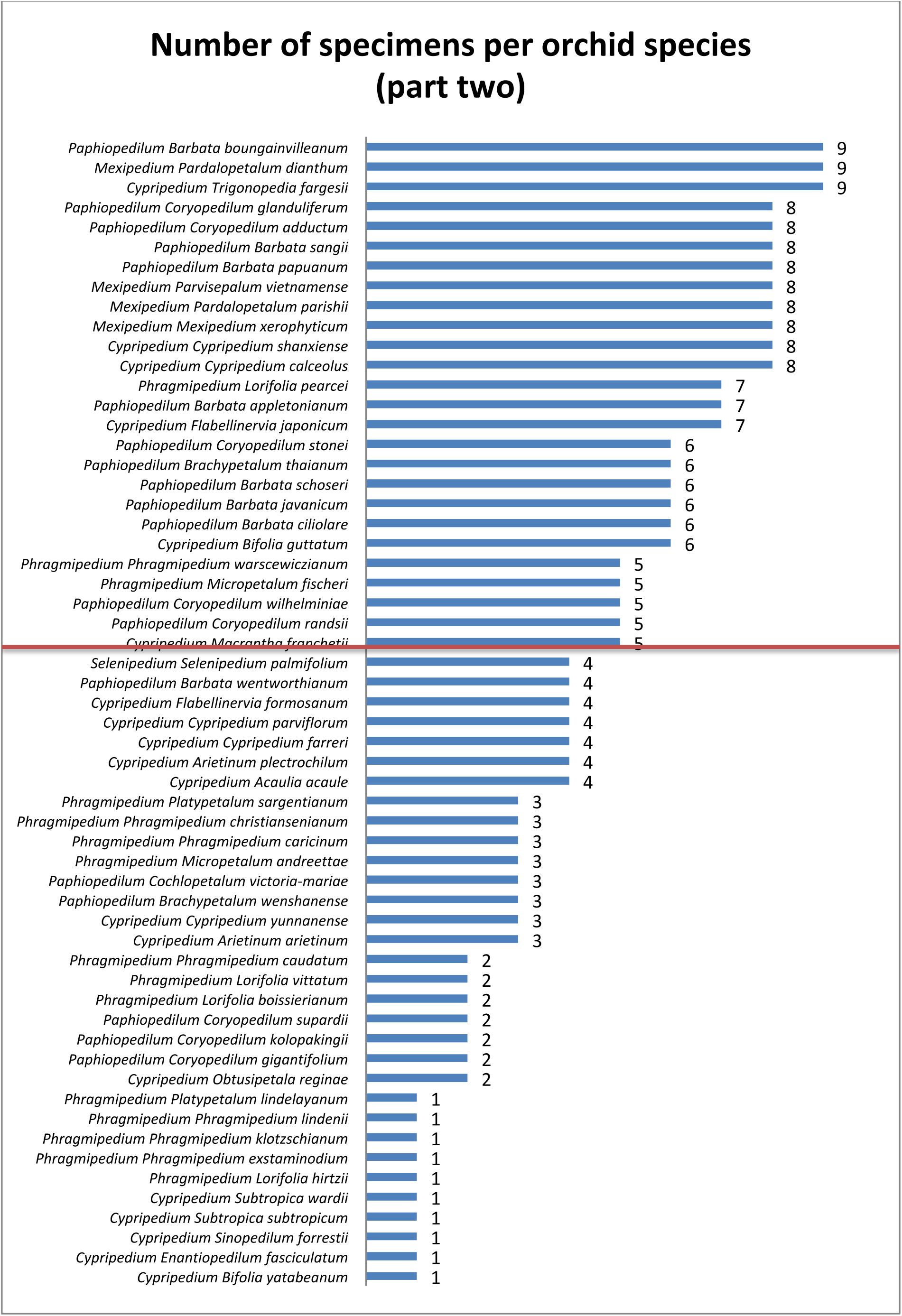
Number of specimens per orchid species part two. The red line indicates the cut-off for the selected dataset (n=5).

### Extracted image features

The ROI that was determined in pixel units was: 0, 900, 2350, 2500. The format is X, Y, Width, Height, where the upper left corner of the picture is pixel 0. This ROI was transformed into fractions of total picture size: 0, 0.3907, 0.215, 0.8467. The format is X1, X2, Y1, Y2, where X1 and Y1 are in the upper left corner, X2 goes to the right and Y2 goes to the bottom. In *Figure 8* the ROI is shown in a red rectangle. All butterfly pictures were cropped to this ROI before proceeding with other steps.

**Figure 8.**
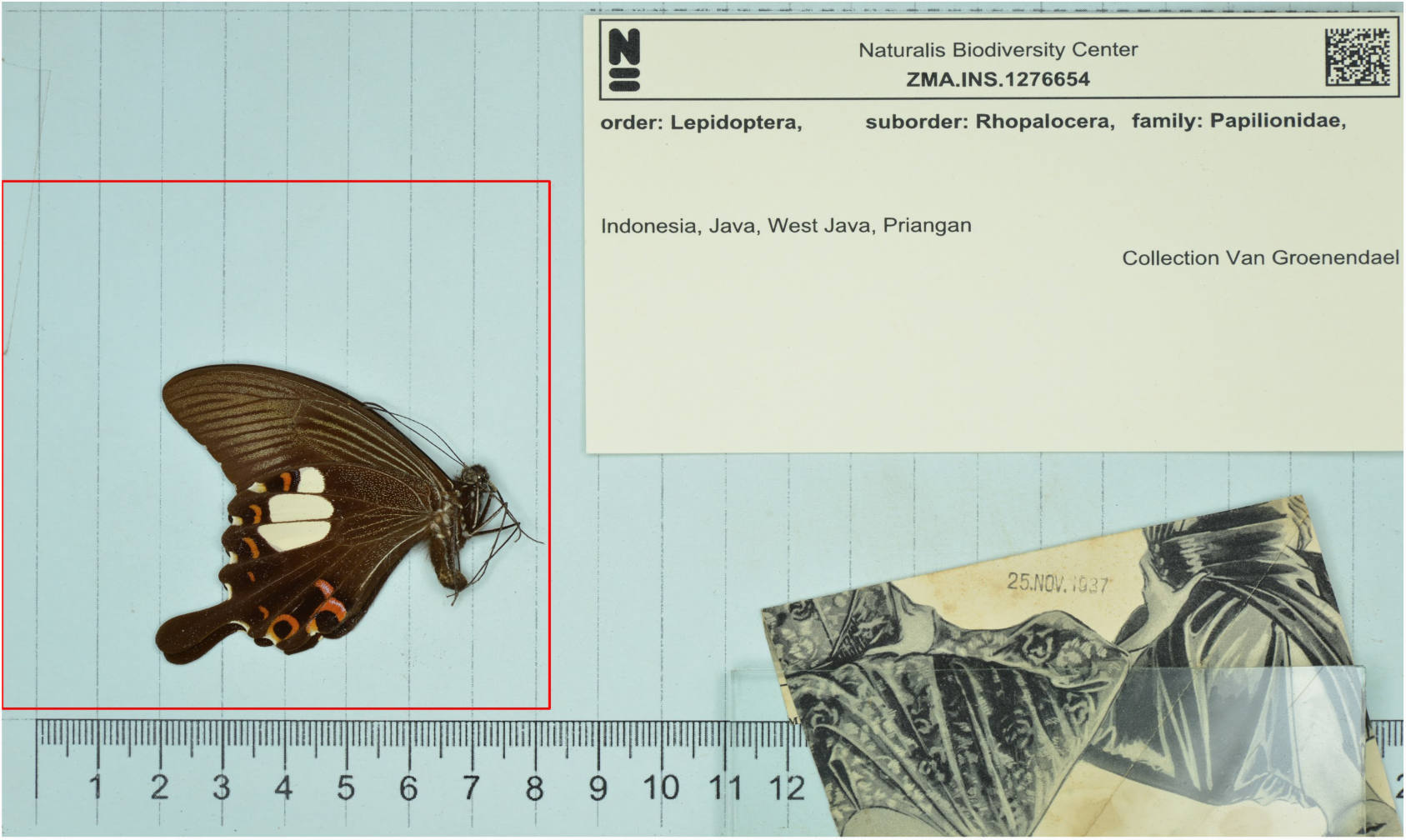
Standardized picture with the Region Of Interest in a red rectangle. All butterflies in the reference dataset are placed inside this region.

Subsequently, the butterfly images were brightness thresholded (truncated) before features were extracted with the SURF algorithm, to avoid the faint lines in the background to be seen as features as well. In *Figure 9*, examples of the detected key points with and without this step are shown. Every blue circle is one detected feature.

**Figure 9.**
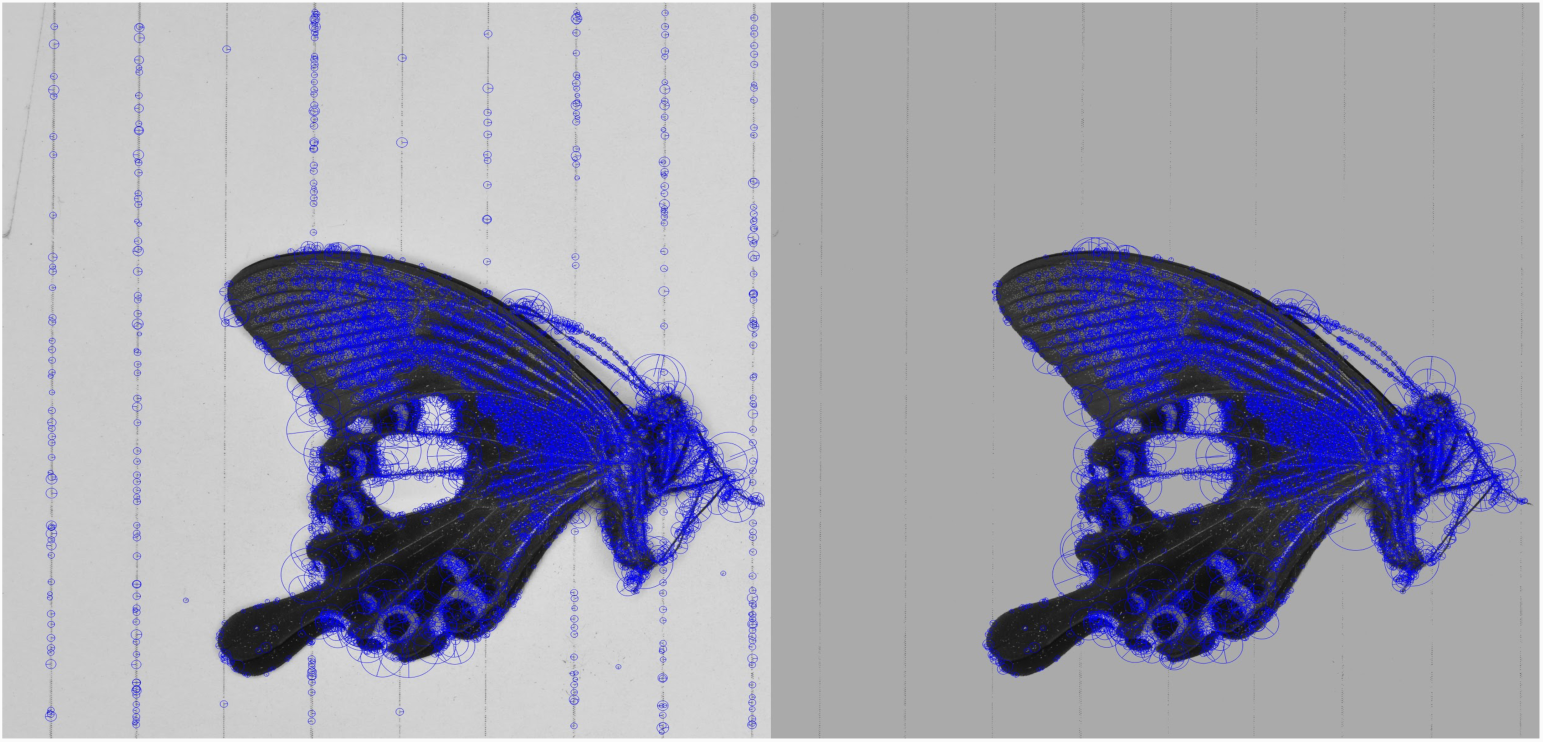
Detected features made visible. On the left the detected features before truncating the image, on the right after brightness thresholding (truncating). On the left side the lines in the background are seen as features as well.

*Table 3* shows results of the nbc-bag-of-words.py script for the different datasets. Three *k* values were applied to the first two butterfly datasets. The results show that the higher the *k* value, the longer it takes for the script to finish. The results also show that the total running time increases with the total number of features. The spreadsheets that came out of this script are stored in the nbclassify-data repository in the archives-butterfly/BagOfWords directory.

**Table 3.** Results of the script nbc-bag-of-words.py for the different datasets. The results show that the more features there are and the higher the k value for the k-means method, the longer it takes for the script to finish.

| Dataset | Total features | $k$ | Total running time |
| --- | --- | --- | --- |
| Dataset1 | 999,877 | 150 | 5 hours 9 minutes |
| Dataset1 | 999,877 | 500 | 17 hours 36 minutes |
| Dataset1 | 999,877 | 999 | 1 day 9 hours |
| Dataset2 | 2,270,433 | 150 | 15 hours 25 minutes |
| Dataset2 | 2,270,433 | 500 | 2 days 2 hours 32 minutes |
| Dataset2 | 2,270,433 | 1,506 | 5 days 6 hours 1 minute |
| Orchids | 627,832 | 125 | 2 hours 47 minutes |

### Image feature performance plots

The first two principal components (PCs) of the BGR histogram of the first butterfly dataset are shown in *Figures 10-11.* Names are used as datapoints in all PCA plots, because colored dots could not be kept apart by the limited number of colors. Genus *Graphium* groups together in the genus-plot, but all other genera are very scattered. The divergence of the species is shown in the species-plot. Most species aren’t nicely grouped together. The plots of PC1 + 3 and PC2 + 3 of the first butterfly dataset and all plots of the second butterfly dataset are shown in *Appendix L*. The results are similar for both datasets.

**Figure 10.**
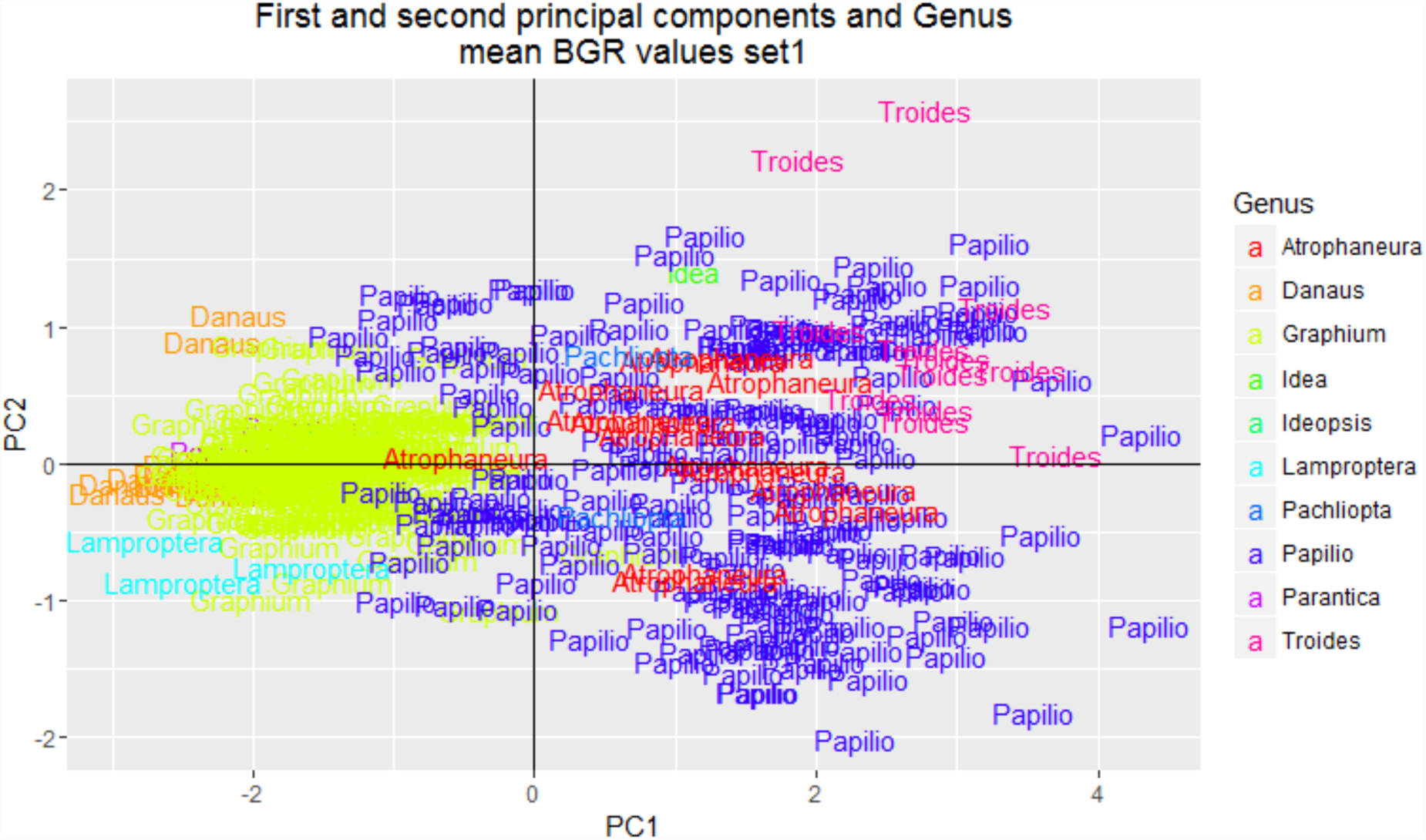
Genus-plot of PC1 and 2 of the mean blue-green-red values of dataset 1. The data points are color-coded on genus and the genus name is used as a label.

In *Figures 12-15* the PCA plots of the first butterfly dataset are shown for the SURF-BOW method. In the first two plots the number of clusters was 150, in the last two 999. The plots of PC1 + 3 and PC2 + 3 of the first dataset and all plots of the second dataset both with different *k*-values are shown in *Appendix M*. The genus *Papilio* is very scattered, but the species-plot shows there are multiple species in this genus that do group together. All other genera and species group together better than with the mean BGR method.

**Figure 11.**
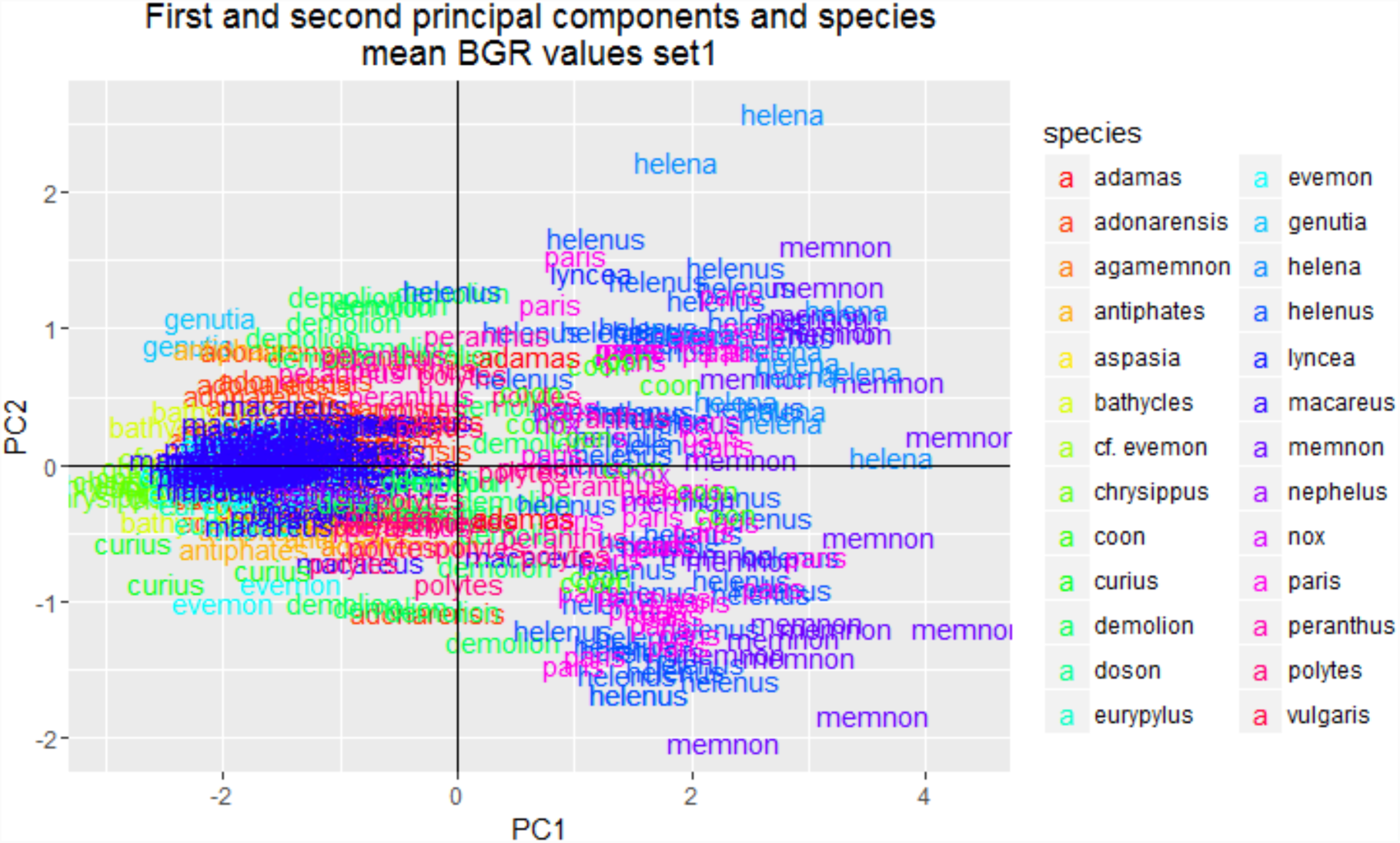
Species-plot of PC1 and 2 of the mean blue-green-red values of dataset 1. The data points are color-coded on species and the species name is used as a label.

**Figure 12.**
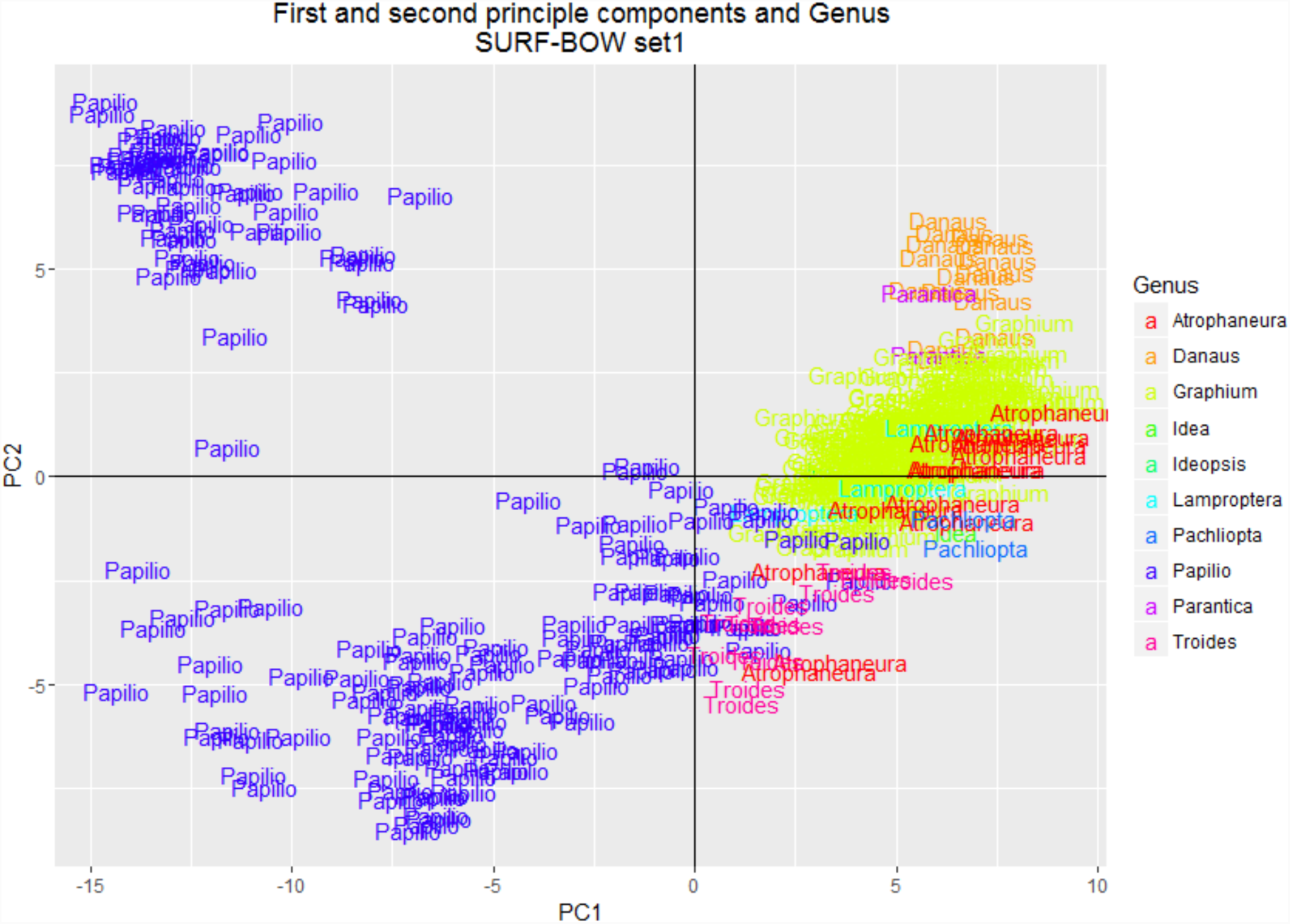
Genus-plot of PC1 and 2 of the SURF-BOW method of dataset 1 with 150 clusters. The data points are color-coded on genus and the genus names are used as label.

*Figures 16-17* show the PCA plots of the BGR-SURF method on the selected butterfly dataset with PC1 + 2. The plots of PC1 + 3 and PC2 + 3 can be seen in *Appendix N*. As with the plots of the SURF-BOW method, the genus *Papilio* is very scattered. The species seem to group together, but the datapoints are very close to each other.

**Figure 13.**
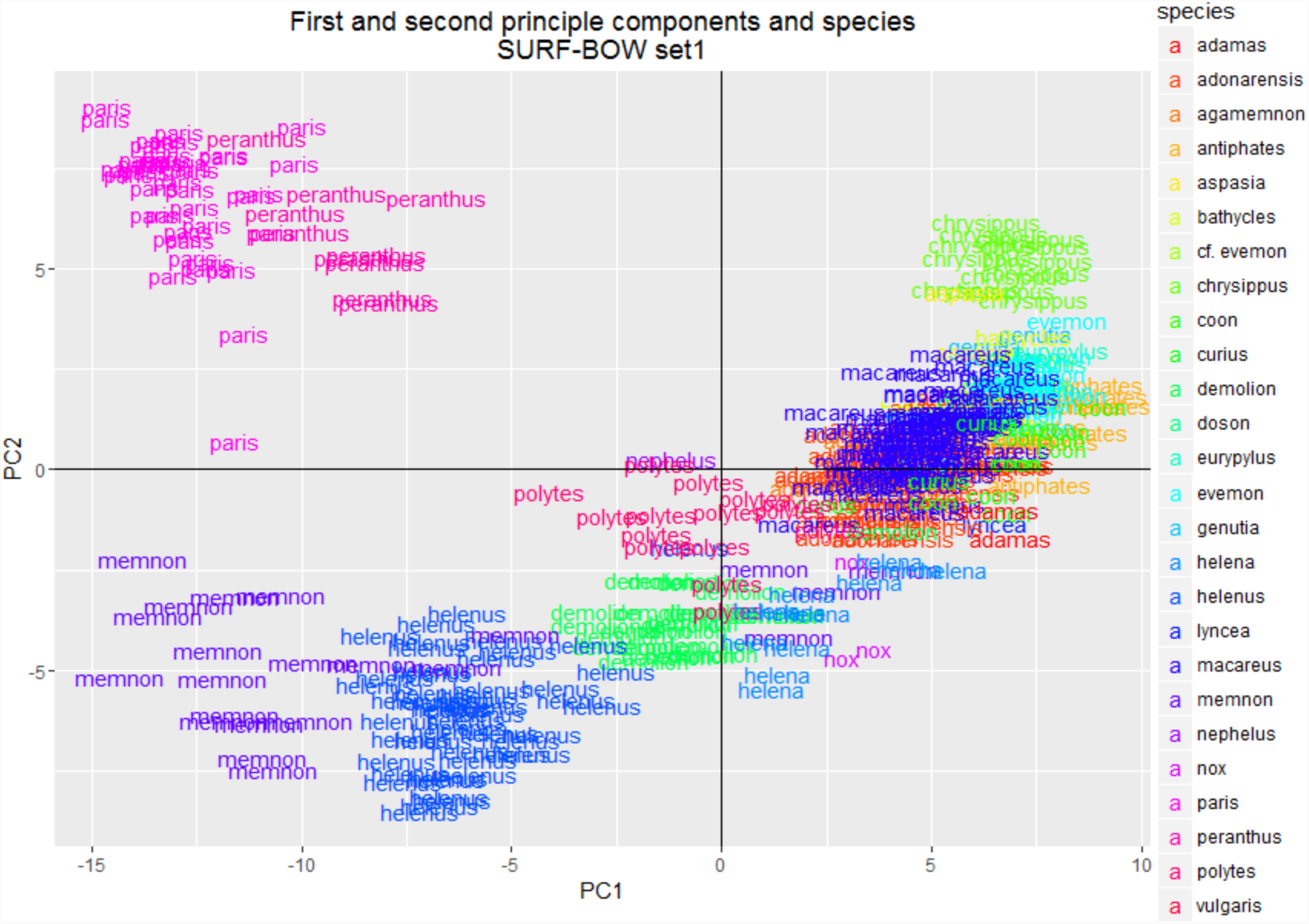
Species-plot of PC1 and 2 of the SURF-BOW method of dataset 1 with 150 clusters. The data points are color-coded on species and the species names are used as label.

**Figure 14.**
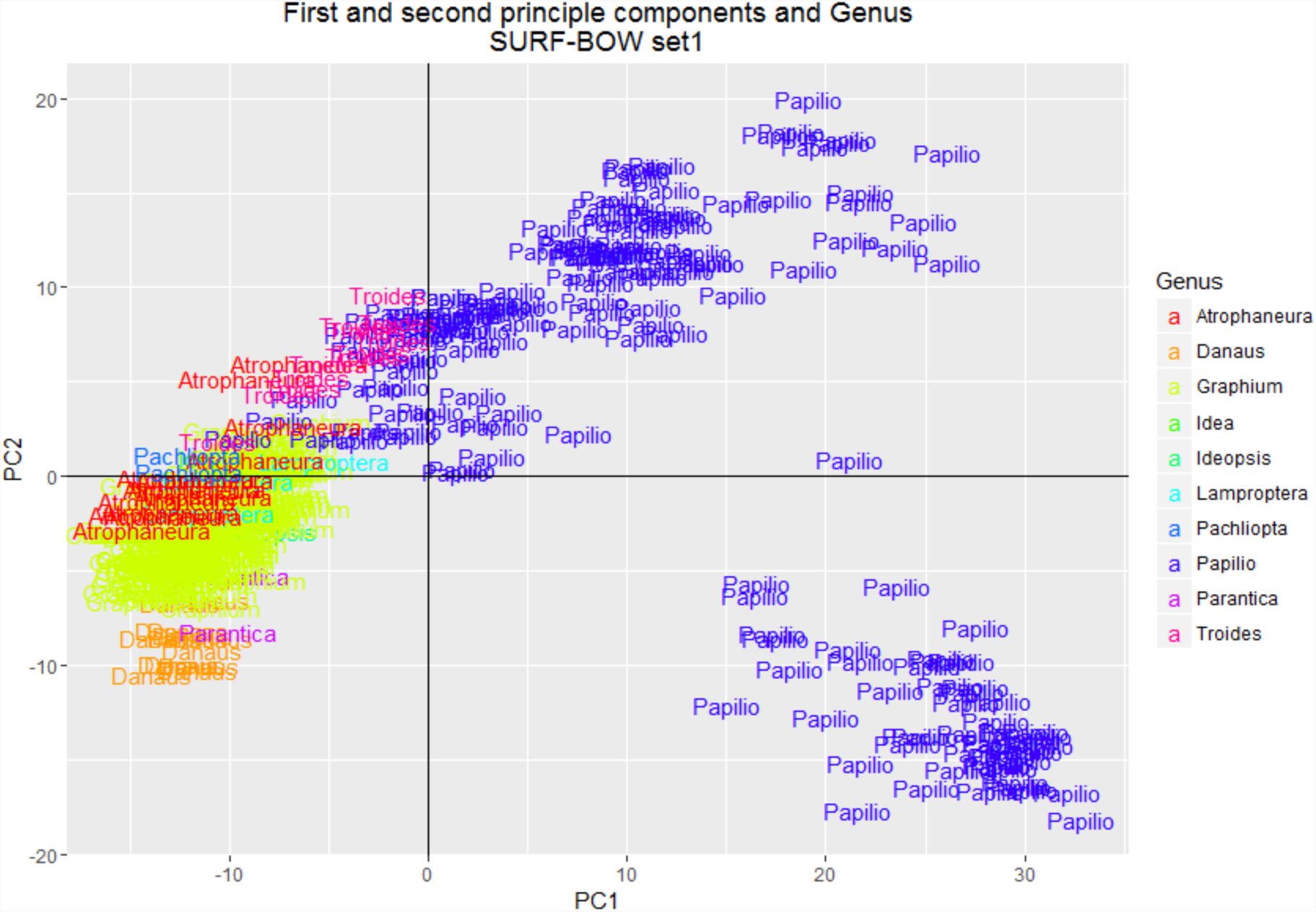
Genus-plot of PC1 and 2 of the SURF-BOW method of dataset 1 with 999 clusters. The data points are color-coded on genus and the genus names are used as label.

**Figure 15.**
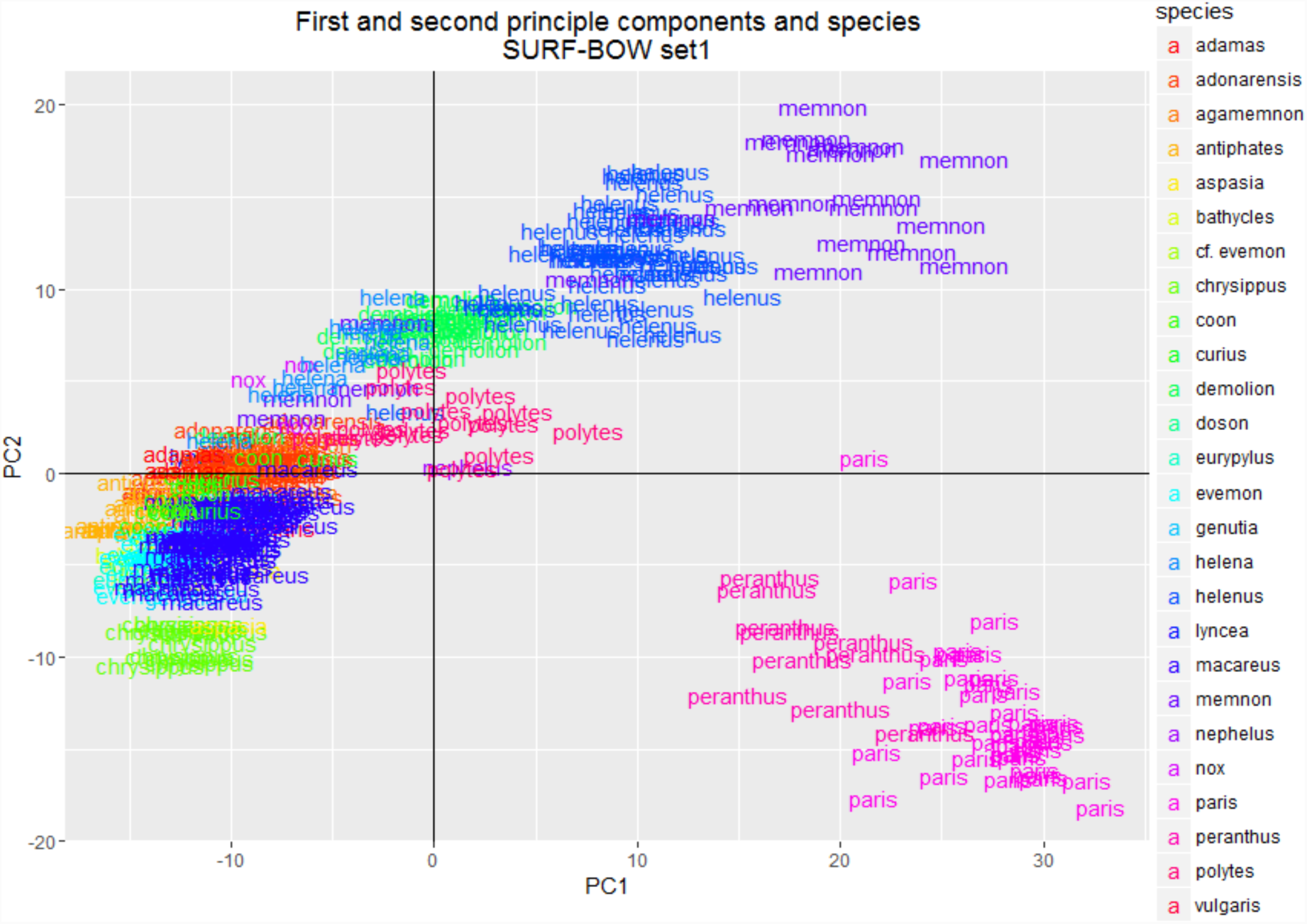
Species-plot of PC1 and 2 of the SURF-BOW method of dataset 1 with 999 clusters. The data points are color-coded on species and the species names are used as label.

**Figure 16.**
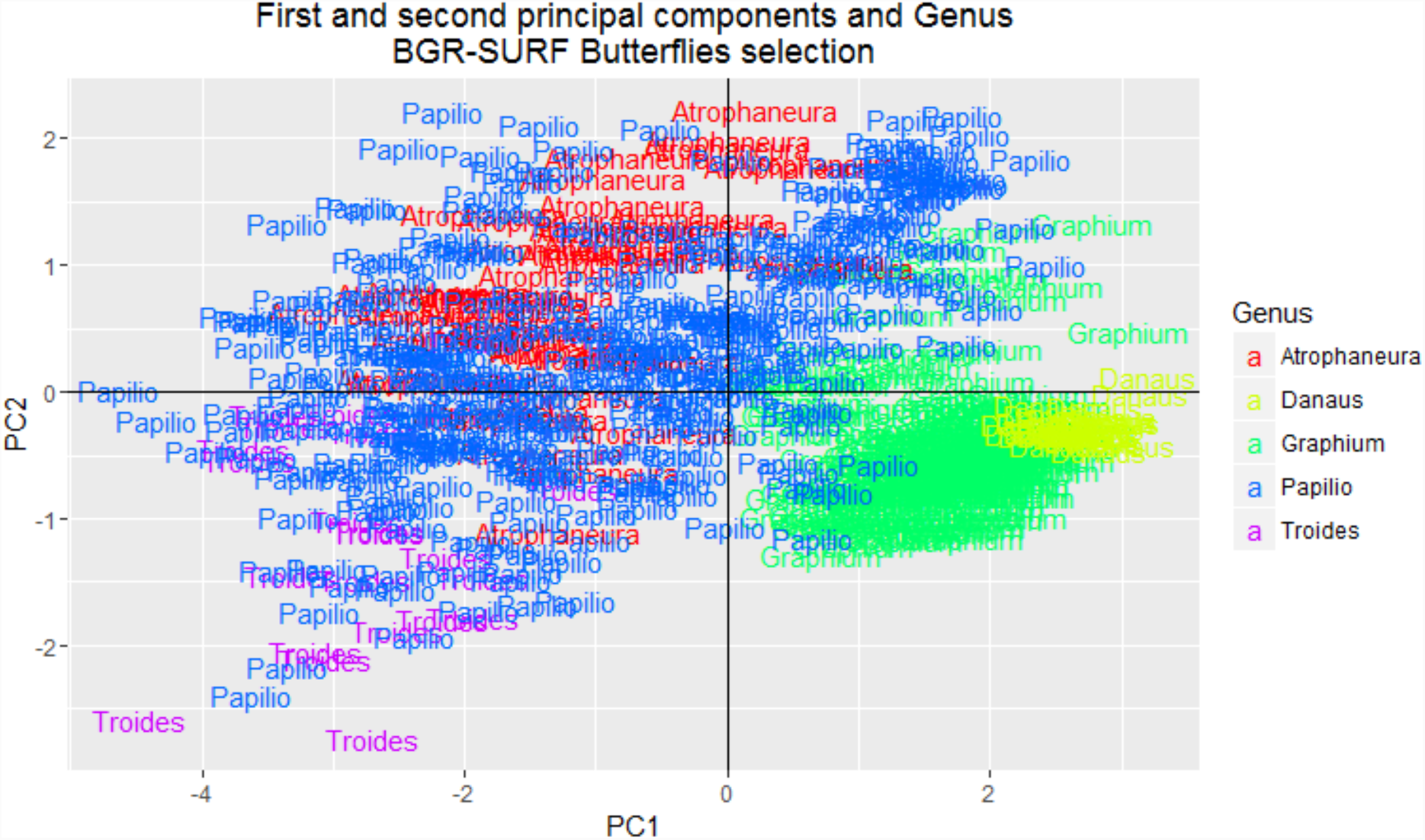
Genus-plot of PC1 and 2 of the BGR-SURF method of the selected butterfly dataset. The data points are color-coded on genus and the genus names are used as label. The SURF-BOW part was done with 150 clusters.

*Figures 18-20* show the PCA plots of the first and second principal components of the orchid dataset. There seems to be some order in the genus-plot, but in the section- and species-plot there is very little order. The other plots can be seen in *Appendix O*.

**Figure 17.**
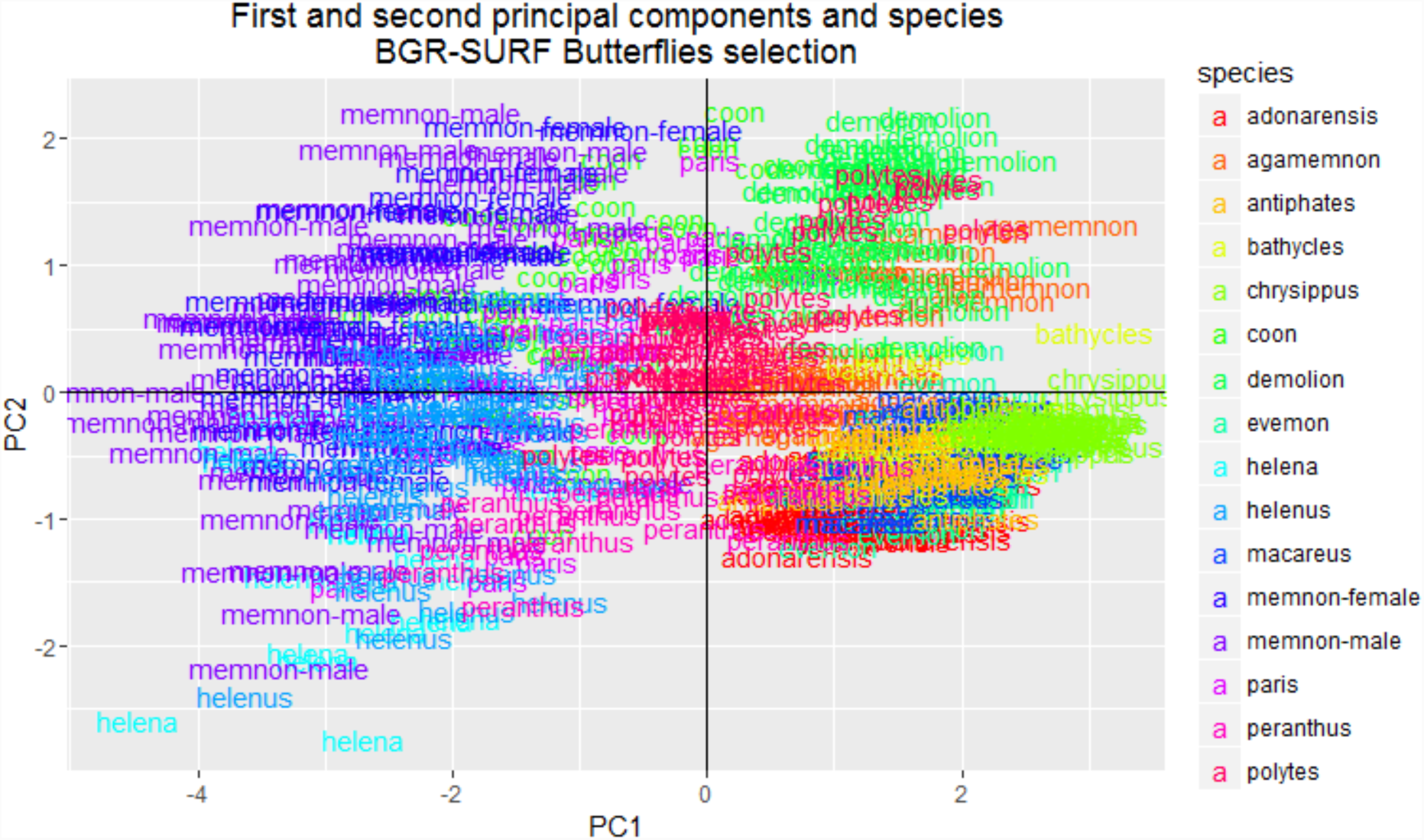
Species plot of PC1 and 2 of the BGR-SURF method of the selected butterfly dataset. The datapoints are color-coded on species and the species names are used as label. The SURF-BOW part was done with 150 clusters.

**Figure 18.**
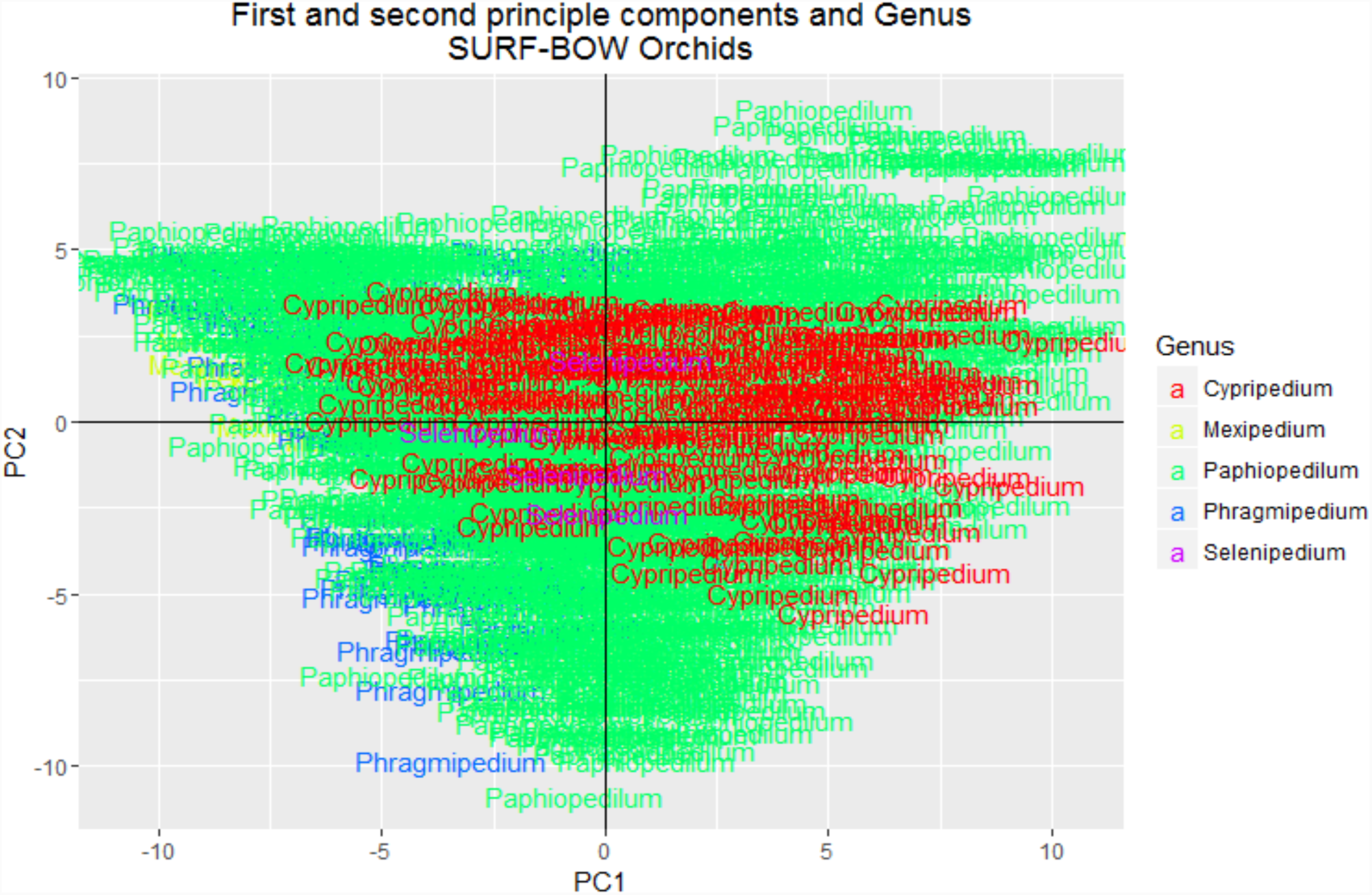
Genus-plot of PC1 and 2 of the SURF-BOW method of the orchid dataset. The data points are color-coded on genus and the genus names are used as label.

### Artificial neural network cross-validation outcomes

*Figures 21* and *22* show the results of the stratified *k*-fold cross validation of the selected butterfly dataset reduced to SURF-BOW features. *Figure 21* displays the results for classification on genus and species separate from each other. In *Figure 22* the results are displayed for the classification on both genus and species. These figures show that most classifications are correct. Less than 5% is classified incorrectly or ambiguously. The rest of the specimens are classified as unknown, which is about 15-20%.

**Figure 19.**
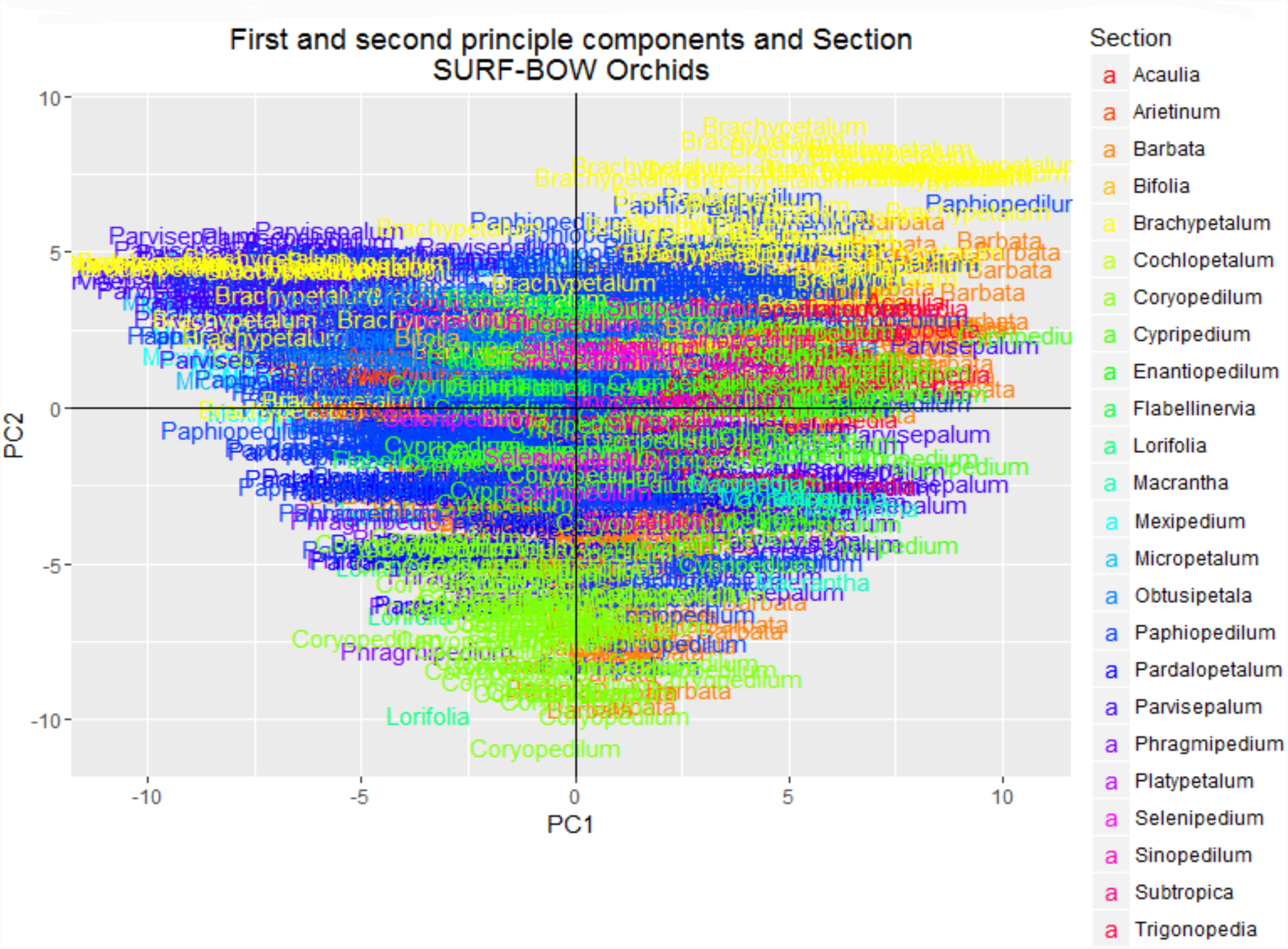
Section-plot of PC1 and 2 of the SURF-BOW method of the orchid dataset. The data points are color-coded on section and the section names are used as label.

**Figure 20.**
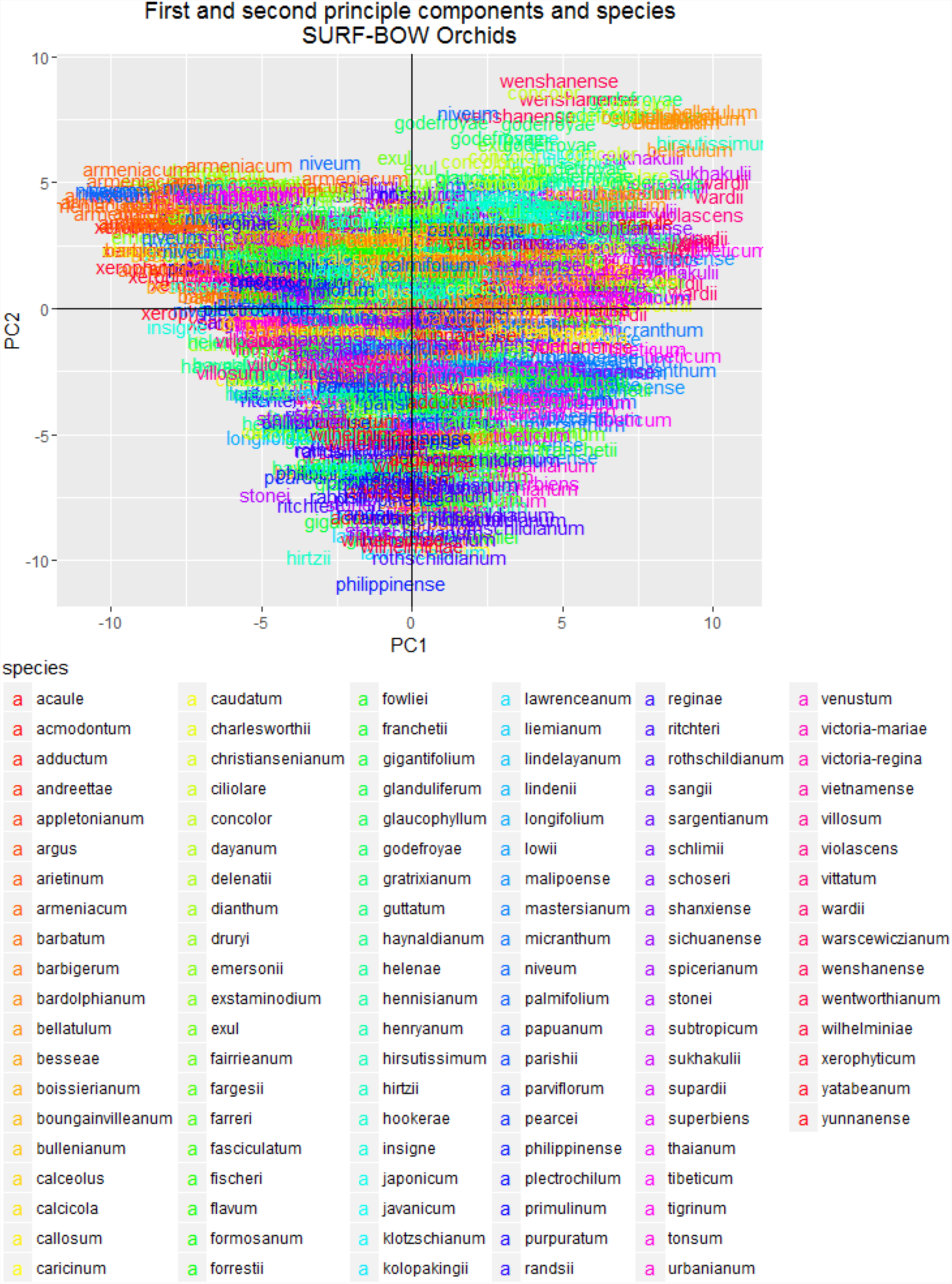
Species-plot of PC1 and 2 of the SURF-BOW method of the orchid dataset. The data points are color-coded on species and the species names are used as label.

**Figure 21.**
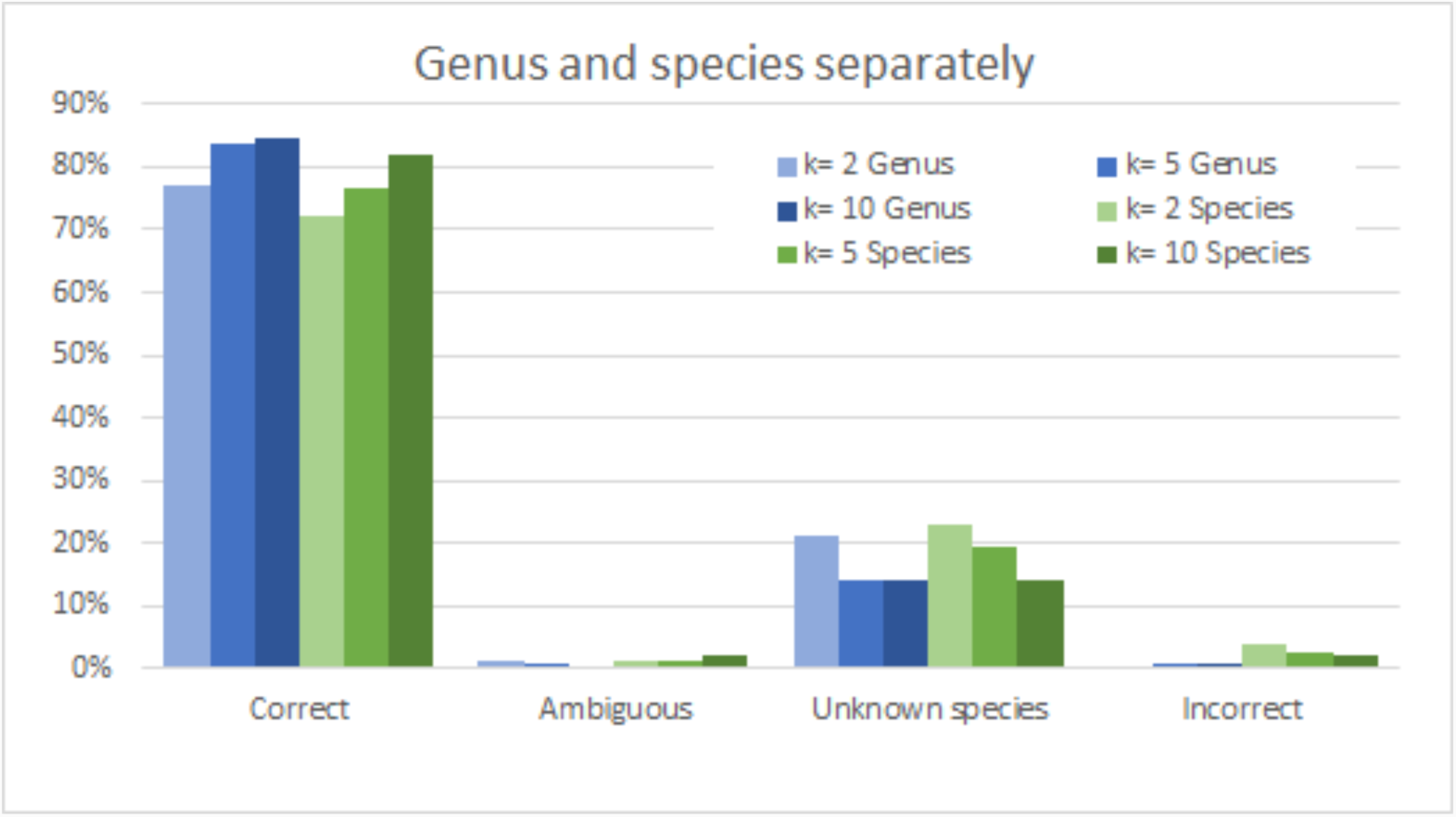
Results of the stratified k-fold cross validation of the selected butterfly dataset with the SURF-BOW method on 150 clusters, classified on genus and species separately. If a prediction was not correct, most of the time the program classified it as unknown.

**Figure 22.**
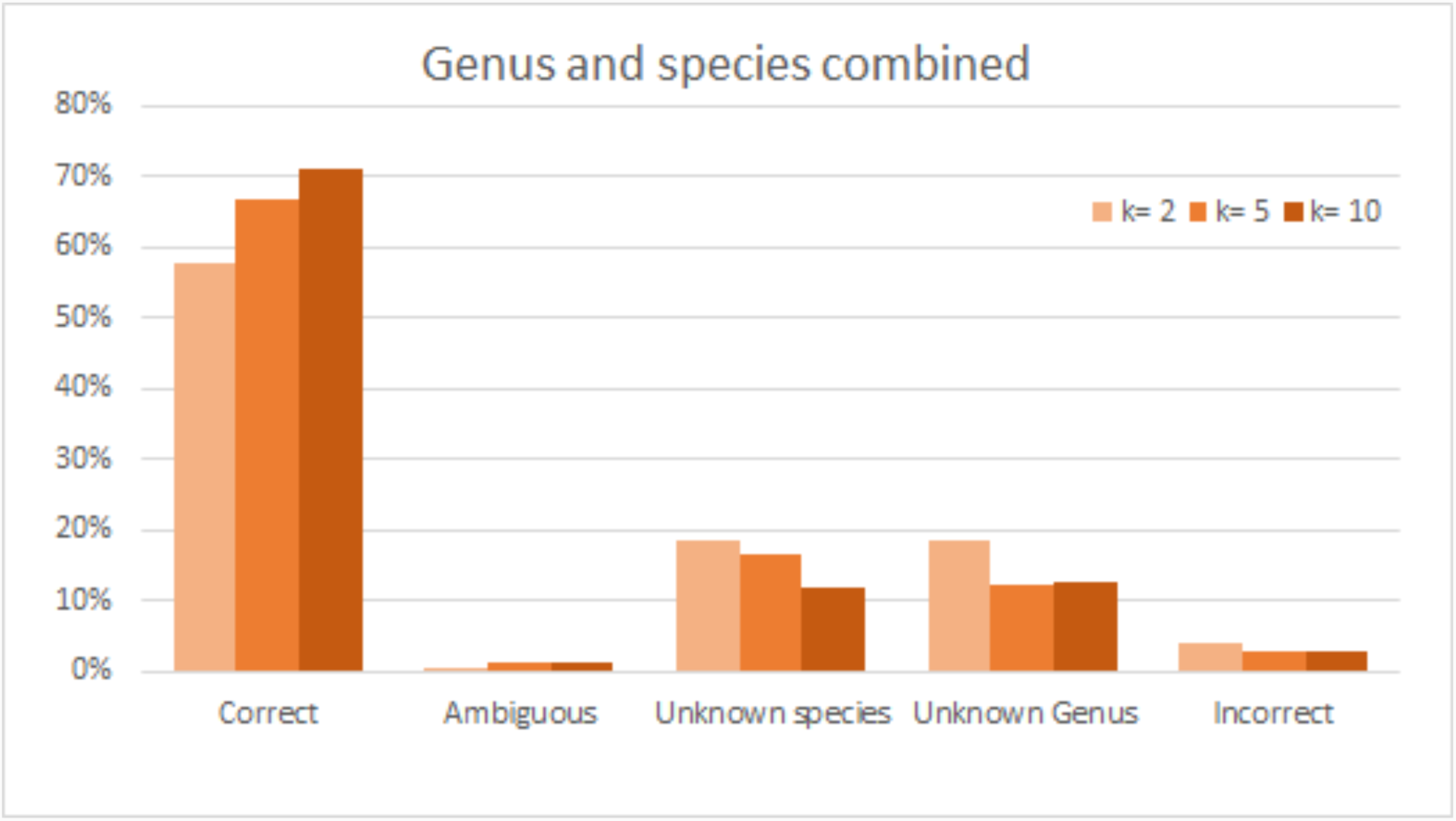
Results of the stratified k-fold cross validation of the selected butterfly dataset with the SURF-BOW method on 150 clusters, classified on both genus and species. Less than 20% had an unknown classification for genus or species. Less than 5% was incorrect or ambiguous.

An extra analysis was done on the selected butterfly dataset for the combined genus-species classification with *k* = 10. The number of individuals in the training dataset is plotted against the percentage correct and incorrect in *Figure 23.* Apart from some exceptions, with fewer than about 15 specimens per species, the percentage correct is lower than 50% and the percentage incorrect is above 10%. However, increasing the number of individuals improves the results (R^2^=0.6326, *p*=4.2459E-07).

**Figure 23.**
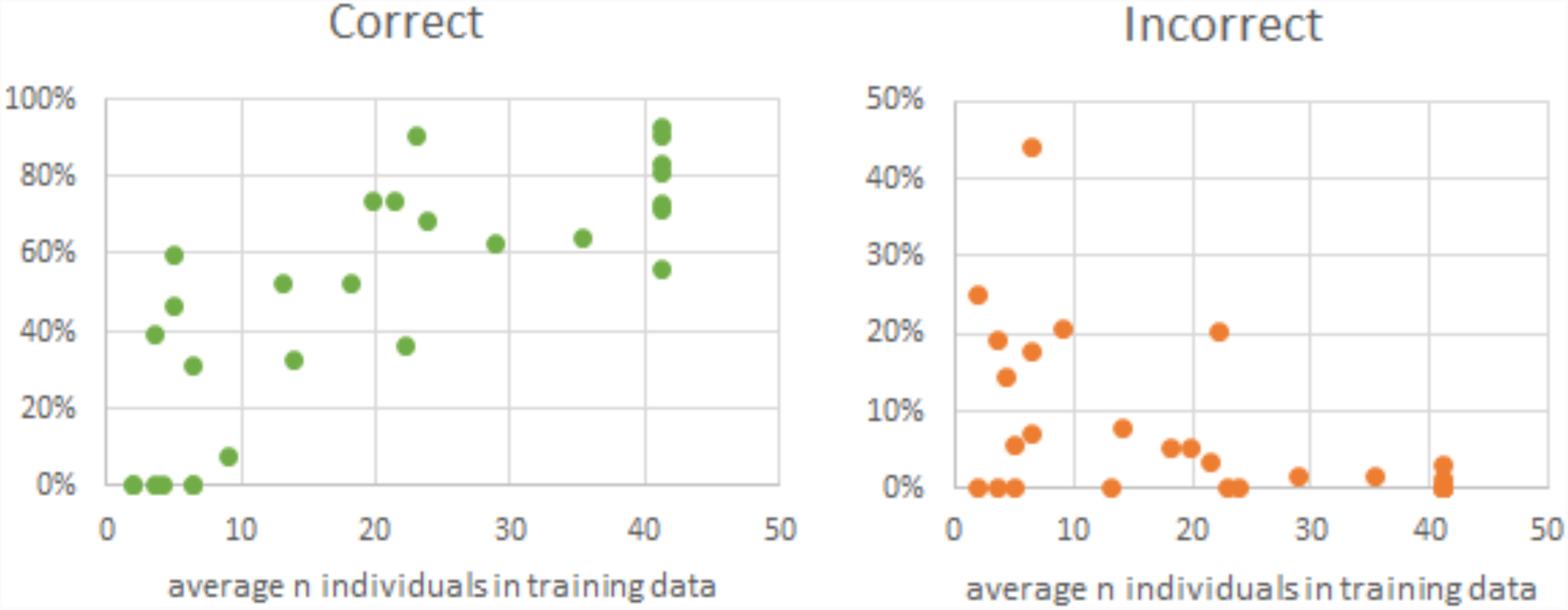
Average number of individuals in training data for the SURF-BOW method for the selected butterfly dataset, classified on both genus and species. Results for k=10. Apart from some exceptions, with less than about 15 specimens per species, the percentage correct is lower than 50% and the percentage incorrect is above 10%.

The SURF-BOW method was also applied to the orchid dataset. The results of the stratified *k-*fold cross validation of the nbc-trainer script on this dataset are displayed in *Figure 24.* The results on *k* = 10 are similar for the three classifications: 24-25%.

**Figure 24.**
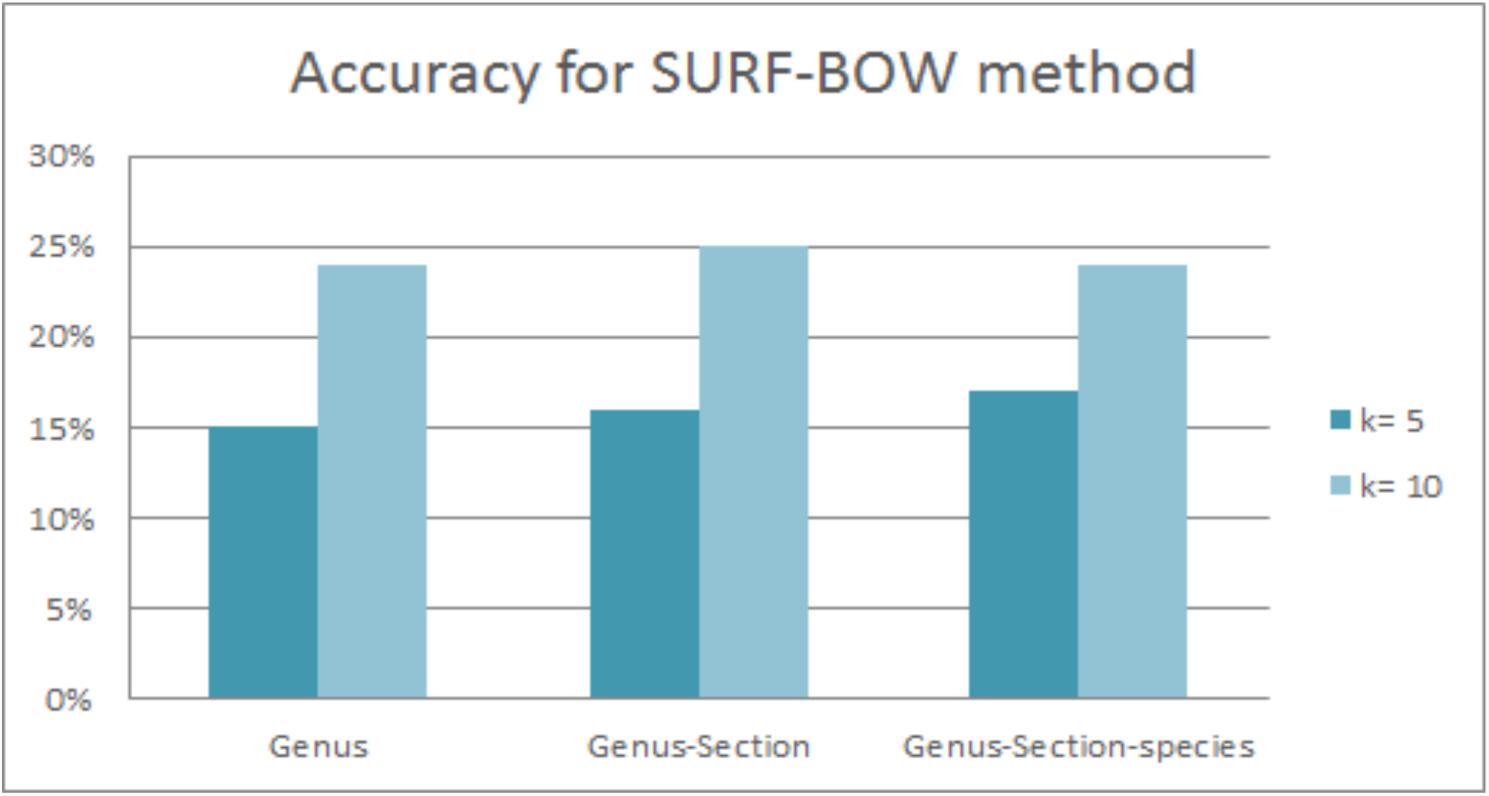
Result of the stratified k-fold cross validation of the SURF-BOW method for the orchid dataset. The best classification had an accuracy of 25%.

In *Table 4* the results of the validation option of the nbc-trainer script are shown for the different feature extraction methods for both orchids and butterflies. A stratified *k*-fold cross validation was performed on the data with different *k*-values. The highest accuracy for the chained prediction of the butterflies with *k* = 10 is 77%, for the orchids it is 28%.

**Table 4.**
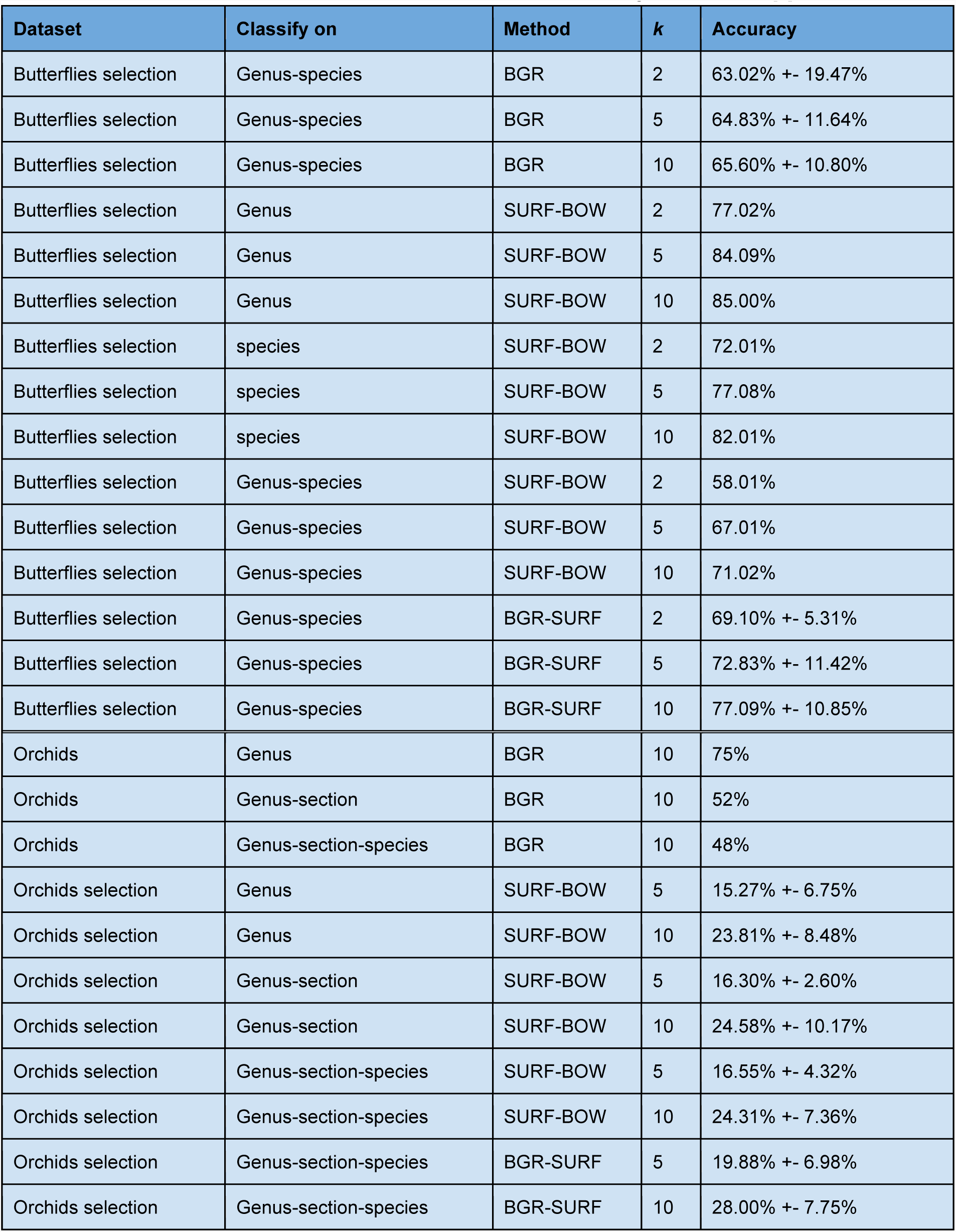
Results of the stratified k-fold cross validation of the artificial neural networks. BGR-SURF is the method where the original method of mean BGR values is combined with the SURF-BOW method. The results from the BGR method for the orchids came from the original framework [3].

| Dataset | Classify on | Method | k | Accuracy |
| --- | --- | --- | --- | --- |
| Butterflies selection | Genus-species | BGR | 2 | 63.02% +- 19.47% |
| Butterflies selection | Genus-species | BGR | 5 | 64.83% +- 11.64% |
| Butterflies selection | Genus-species | BGR | 10 | 65.60% +- 10.80% |
| Butterflies selection | Genus | SURF-BOW | 2 | 77.02% |
| Butterflies selection | Genus | SURF-BOW | 5 | 84.09% |
| Butterflies selection | Genus | SURF-BOW | 10 | 85.00% |
| Butterflies selection | species | SURF-BOW | 2 | 72.01% |
| Butterflies selection | species | SURF-BOW | 5 | 77.08% |
| Butterflies selection | species | SURF-BOW | 10 | 82.01% |
| Butterflies selection | Genus-species | SURF-BOW | 2 | 58.01% |
| Butterflies selection | Genus-species | SURF-BOW | 5 | 67.01% |
| Butterflies selection | Genus-species | SURF-BOW | 10 | 71.02% |
| Butterflies selection | Genus-species | BGR-SURF | 2 | 69.10% +- 5.31% |
| Butterflies selection | Genus-species | BGR-SURF | 5 | 72.83% +- 11.42% |
| Butterflies selection | Genus-species | BGR-SURF | 10 | 77.09% +- 10.85% |
| Orchids | Genus | BGR | 10 | 75% |
| Orchids | Genus-section | BGR | 10 | 52% |
| Orchids | Genus-section-species | BGR | 10 | 48% |
| Orchids selection | Genus | SURF-BOW | 5 | 15.27% +- 6.75% |
| Orchids selection | Genus | SURF-BOW | 10 | 23.81% +- 8.48% |
| Orchids selection | Genus-section | SURF-BOW | 5 | 16.30% +- 2.60% |
| Orchids selection | Genus-section | SURF-BOW | 10 | 24.58% +- 10.17% |
| Orchids selection | Genus-section-species | SURF-BOW | 5 | 16.55% +- 4.32% |
| Orchids selection | Genus-section-species | SURF-BOW | 10 | 24.31% +- 7.36% |
| Orchids selection | Genus-section-species | BGR-SURF | 5 | 19.88% +- 6.98% |
| Orchids selection | Genus-section-species | BGR-SURF | 10 | 28.00% +- 7.75% |

### Version control

Imgpheno repository:

Changes were made to the __init__.py script.

nbclassify repository:

Two scripts were changed and four scripts were added to the nbclassify/nbclassify/scripts directory.

Six scripts were changed in the nbclassify/nbclassify/nbclassify directory.

nbclassify-data repository:

The directory ‘archive-butterflies’ was added with all its content.

## Discussion

### Image data and metadata

When the pictures were tagged on the Flickr website, it was expected that the number of tagged images would be the same as the number of specimens in the dataset. For butterfly dataset two and three this was the case, but dataset one tagged 383 images, while there were only 382 specimens in the dataset. It appeared that the image with registration number 1276325 was uploaded twice to the Flickr website. One of these was removed from the Flickr website before proceeding with the other steps.

In the selected butterfly dataset, *Papilio polytes* was first split in two groups according to sex. After the first validation of the artificial neural network, it appeared that most of the times the *P. polytes* males were classified as females and the other way around. *P. polytes* is known for its mimicry in the female specimen [17, 30] and has two types of phenotype. It therefore was decided to not split this species into sex. All 27 female specimens were used in the definitive dataset and males were added for a total number of 50. The three butterfly datasets contain 1845 pictures, divided over 48 species in 20 genera. The selected dataset had 732 pictures of 33 species in 10 genera. The 617 pictures that form the definitive dataset contain 17 species and 5 genera.

Families *Pieridae* and *Riodinidae* and some genera and species of the other families had too few specimens to end up in the selected dataset and/or the definitive dataset. Therefore specimens of these groups could not be recognized by the ANN, and would be classified as either unknown or incorrect. Since the number of specimens of these groups in the total reference dataset was so limited, it is expected their total number in the whole collection will also be limited and could therefore be classified manually, assuming the bulk of the collection is classified automatically. If in the future more data becomes available to use in the reference dataset, for example when the classification is done on (a part of) the whole collection, these species might have enough specimens to add to the system, so they can be classified as well.

A selection of the orchid dataset was made with at least five specimens per species. The results of the butterflies show that for the SURF-BOW method upward of 15 specimens per species are necessary (and the more the better) for the reference dataset in order to get a good prediction, see *Figure 23*. Since the number of specimens per species in the orchid dataset is so limited, it was decided not to apply the minimum of 15 to the orchids. If in the future more orchid pictures become available to use as a reference, the minimum of 15 can be applied and the method can be checked to be more accurate.

### Image feature extraction

The pictures of the butterflies were made in standardized position, but when the ROI was determined it turned out that not all the pictures had the same resolution. The explanation for this is that when the first pictures were taken, the best setup still needed to be found. Only the first 70 pictures that were taken have a size of 1920 x 1282 pixels (width x height). Also instead of a printed registration number and QR code on the card, they had a sticker in the top right corner of the card with this information. These pictures were taken in November 2015. The size of the pictures was changed to 6016 x 4016 pixels in December 2015, to improve the quality of the pictures. In addition, from that moment on, all the registration numbers and QR codes were printed on cards. In March 2016 the size was changed once again, this time to 4256 x 2832 pixels. This was done to limit the storage space. The pictures of 1920 x 1282 pixels had an average size of 200kB. The pictures with 6016 x 4016 pixels had an average size of 9.04MB and the pictures that had 4285 x 2832 pixels had an average size of 4.62MB.

The *k*-means clustering for the BagOfWords method needs a lot of RAM. With a small subset of the datasets it was possible to run the script on a local Windows computer (4GB RAM, dual core, 1.50GHz/core), but with the whole first butterfly dataset it ran into memory constraints. It therefore was necessary to perform all the runs of this method on a cloud instance (64GB RAM, quad core, 2.60GHz/core). All other steps of the process can be performed with the whole datasets on a local Windows computer, but on the cloud it will go much faster.

### Image feature performance visually inspected

There is little difference between the clustering of genera and species when the butterfly PCA plots of 150 and 999 clusters are compared. *Table 3* showed a large difference in running time between the two different *k*-values. Therefore a *k* of 150 was found to be better.

To make the results of the PCA visible, three 2D plots were made to get an idea of the 3D distribution of the data points. It would probably be more efficient to have a 3D plot that could be rotated by the user. This type of plot was also created, but this, however, could not be saved in a useful way. Therefore it was decided to work with the three 2D plots. If in the future it is possible to save a 3D plot in a way the user can rotate it after it has been saved, this will be the favoured way to display these results.

The PCA plots for the SURF-BOW method and BGR-SURF method of the butterflies show a clustering of the species, but the genera these species belong to are more widely scattered. A reason for this is that the clustering was done based on image features, in this case wing patterns. Presumably this is because the taxonomic classification was not done based on morphological similarity (i.e. phenetics), but based on evolutionary relatedness, and close relatedness does not necessarily imply great similarity in wing patterns [31]. Therefore is it normal to see a more diverse genus plot and obvious groups in the species plot.

For the orchids, the PCA plots seem to have less structure. However, the number of species (or sections) is so large and the number of specimens per species is so low, that there is not enough data to make visible clusters. Therefore it was decided to create an ANN and perform the validation on it, even though the PCA indicated that it would be a bad method for this dataset.

### Artificial neural networks

As *Figures 21-22* describe, with the SURF-BOW method in about 85% of the cases the genus was predicted correctly for the butterflies. In 71% of the cases, both genus and species were correct. The combined BGR-SURF method seems to do slightly better for both genus and species prediction. This is probably because the SURF algorithm detects the patterns on the wing, and the BGR method adds information about the color distribution over the bins, where information about the shape of the butterfly comes from, like a tail on the wing. The combination of these methods creates more specific information to classify on, which leads to greater accuracy of the ANN. The system classified most specimens it was uncertain about as ‘unknown’, so only 1-4% was incorrect or ambiguous, see *Figures 21-22*. This is the preferred way, because an unknown specimen can be inspected visually, while a false positive might go undetected.

In the original OrchID framework, the accuracy of the system was 75% on genus, 52% on genus and section and 48% on genus, section and species. For the orchids, the new SURF-BOW method performs much worse than the original mean BGR method: less than 25% accurate for any of the three ranks. The combined method of mean BGR and SURF-BOW is even worse for the orchids. A reason for this might be that the SURF method uses changes in pixel intensity to detect a feature in an image, like the patterns on a butterfly wing. However, unlike the butterflies, these orchids do not have patterning in fixed locations on their surface to extract features from, what makes it hard to identify them with this method. The SURF features are apparently not discriminating enough for this dataset and when added to the mean BGR values, the SURF features perhaps swamp those extracted with the BGR method.

*Table 4, Figures 21-22* and *Figure 24* show the higher the *k* value, the better the accuracy gets. An explanation is that the higher the *k* value, the more specimens are in the training dataset, so there is a larger sample to generalize from and the risk of overfitting is reduced. This leads to a better recognition of a species than with fewer specimens to train on. *Figure 23* shows the same: the more specimens in the training dataset, the better the results. At least 15 specimens per species are necessary in the training dataset to get good results, so the definitive butterfly dataset is made with at least this number of specimens per species. The graph shows that the results improve with more than 15 specimens per species and it suggests that the improvements continue with over 40 specimens. When more data is available to use in the training dataset, this can be investigated. For the orchids, in the future if there is more reference data available, the same criteria can be applied to the dataset and checked to perform better.

### Version control

At the start of the project it took a long time to figure out where everything was stored on the GitHub repositories. There are so many directories with subdirectories that have the same name, the overview gets lost very quickly. It would be nice if this could be improved, but since all scripts are dependent on each other, this is a tricky job to do. Maybe in the future this can be done. For now all original structure is kept.

## Conclusions

When looked at the PCA of the mean BGR values and the SURF-BOW method, the new method works better for the butterfly pictures. Given the accuracy of the system, the combination of both methods, i.e. BGR-SURF, is the best of the tested methods to classify the pictures of the Javanese butterfly collection. The accuracy may be improved by adding more data to train the network with, so there is a larger sample to generalize from and the risk of overfitting is reduced. This leads to a better recognition of a species than with fewer specimens to train on.

Based on the accuracy of the artificial neural networks for the orchids, it is clear that the original method worked better to classify this group of slipper orchids. However, there are very few specimens per species in the dataset and a large number of species. If there were more specimens per species in this dataset, more variance could be overcome and a better classification system could be made. The SURF-BOW method does not improve the classification of slipper orchids compared to the original BGR method, and neither does BGR-SURF.

### Future work

The classification for the whole Javanese butterfly dataset must still be run with the ANN of the definitive dataset. When (a part of) this collection has been classified, the reference dataset can be extended with the classified specimens, so more specimens per species will be available and the system will get more accurate. The specimens that are classified as unknown may be looked at by an expert. If the number of specimens is at least 15, more species can be added to the dataset to create better ANNs. Also species that now had only one sex in the dataset can be checked for dimorphism to decide if they should be treated as separate species as well.

The used methods can be tested with other datasets, like mosquitoes, shells, beetles, etcetera. As for the orchids and butterflies, different datasets require different methods, so it is possible that the mean BGR-values and/or SURF-BOW method will not work for other cases. This has to be investigated in the future.

The classification is done by an artificial neural network at every level of the hierarchical taxonomy. However, a common approach is to have multiple ANNs, in a committee, decide on a classification. Also, in the future other machine learning algorithms can be tested to see if there are better systems, for example support vector machines. In addition, other image feature algorithms can be explored, such as CNN, HOGS, and HoCS.

Going forward, several steps need to be taken to consolidate the code base. For example, as stated before, two versions of the FlickrAPI are now needed to connect to the website. The connection method ‘authenticate via token’ used in nbc-harvest-images needs to be changed to the method ‘authenticate via browser’, so it can use the latest version of the Flickr API as well. In addition, image management on Flickr needs improvement, in particular in handling image duplicates and in managing image sets for separate projects.

It would be good if the nbclassify repository was split into two projects. However, this has to be done very carefully, since all scripts are dependent on each other. Lastly, the data repository must be cleaned up to present the results more clearly (in the form of supplementary data to a publication), and several code bases (imgpheno and the two parts of nbclassify) must be uploaded to PyPI. These and other milestones are added to the GitHub repositories as issues and can be closed once the task is fulfilled.

## Abbreviations and glossary

ANN: Artificial Neural Network A machine learning technique that mimics the brain: input signals enter at input nodes, they are weighted and transformed according to their importance along nodes in one or more ‘hidden layers’ and combined together to create output signals at the output nodes. In this project the input signals will be the image features, the output signal is the classification.
API: Application Programming Interface A set of definitions that makes it possible for software programs to easily interact and exchange information with each other, without the necessity to know the exact functionalities of the other program.
BGR: Blue-Green-Red Refers to the method where an image is divided into bins and the mean BGR values of each bin are calculated and used for the image analysis.
BGR-SURF: Combination of the BGR method and the SURF-BOW method.
BOW: Bag-Of-Words Clustering method, explained in detail in *Appendix I*.
CSV: Comma-Separated Values Tabular text file format with a comma as field separator.
descriptor: A vector of 128 elements to describe a feature and its surrounding area, explained in detail in *Appendix A.*
FANN: Fast Artificial Neural Network Library Open source library to create and use ANNs.
features: Characteristics of an image, like corners and edges of stripes and spots. A feature will be described by a key point and a descriptor.
key point: The coordinate of a feature in an image.
md5sum: Digital checksum, or “fingerprint”, of a chunk of data or a file.
papillotte: Little envelope that a butterfly specimen is stored in.
PC: Principal Component
PCA: Principal Component Analysis
ROI: Region Of Interest
SURF: Speeded Up Robust Features Algorithm that detects steep changes in pixel intensity to locate features in an image.
SURF-BOW: Combination of the BOW method and the SURF algorithm. The BOW method is used to cluster the output of the SURF algorithm for the image analysis.
TSV: Tab-Separated Values Like CSV, but with a tab as field separator.

